# Conversion of a Viral Glycan Shield into a Binding Anchor via Causal-Driven Antibody Optimization

**DOI:** 10.64898/2026.06.25.734658

**Authors:** Xin Wang, Guanying Zhang, Mingda Hu, Jin Han, Chao Shang, Lejin Zhang, Zhengshan Chen, Ping Huang, Wentao Wang, Xinru Zhao, Yunzhu Dong, Yunxiang Zhao, Peng Lv, Xiaodong Zai, Ruochun Jin, Haotian Wang, Congwen Wei, Xiao Li, Liming Yan, Zhiyong Lou, Hongguang Ren, Junjie Xu, Xiangyang Chi

## Abstract

Therapeutic antibodies are challenged by rapidly evolving pathogens that exploit glycosylation to shield epitopes. SARS-CoV-2 JN.1 exemplifies this, escaping antibodies through the N354-linked glycan. However, targeting glycosylated epitopes remains vacant, as scarce and heterogeneous glycan structures render existing approaches ineffective. Here, we introduce the Antibody Evolution Nexus with Causal-Driven Simulation (AENCS), integrating molecular simulation with causal inference. Applying AENCS to restore S309 efficacy against JN.1, we identified ACC01, exhibiting ∼24-fold improved neutralization. With limited prior knowledge of the N354 glycosylation site, ACC01 stabilized this glycan conformation, facilitating the determination of its cryo-EM structure. Causal dissection revealed how this glycan shield is functionally inverted into a binding anchor through multi-layered interactions. This mechanistic conversion, combined with the conservation of N354 glycosylation, enabled ACC01 to maintain potent activity against the latest variant NB.1.8.1. Collectively, AENCS demonstrates causal-driven antibody engineering can illuminate cryptic glycosylated epitopes, providing viable paradigms for exploring this vacant frontier.

---

Therapeutic monoclonal antibodies are the pillar of contemporary biomedicine, yet the relentless antigenic evolution of pathogens continually erodes their clinical durability [1]. Glycosylation represents a pivotal strategy by which pathogens achieve immune evasion: viruses introduce or remodel N-linked glycosylation sites in the proximity of key epitopes, deploying dynamic steric shields that obstruct antibody recognition [2–4]. Unlike protein sequence mutations, however, glycosylation is a post-translational modification marked by pronounced conformational heterogeneity, making it exceedingly difficult to capture structurally and to target through rational antibody design. Glycoform heterogeneity further confounds structure-based design, as no single static structure represents the native ensemble. Consequently, although glycosylation is pervasive in immune evasion, the design of antibodies targeting glycosylated epitopes remains a largely unexplored frontier.

The SARS-CoV-2 Omicron sub-lineage JN.1 starkly illustrates this challenge [4–7]. JN.1 combines RBD point substitutions with a remodeled N354-linked glycosylation site, and the resultant glycan acts as a dynamic steric shield that directly occludes the conserved core epitope of the S309 antibody [8], substantially ablating its potency [3, 8–10]. The N354 glycan thus serves as a litmus test: it represents a class of glycosylated epitopes that are both functionally decisive and mechanistically elusive, against which existing antibody optimization methods are nearly powerless.

Current antibody optimization strategies fall into two categories: directed evolution via wet-laboratory display technologies [11–14], and computational methods such as protein language models [14–19]. While effective when abundant structural data are available, these approaches are fundamentally incapacitated when confronting glycosylated epitopes. The N354 glycan has not yet been structurally resolved, and its conformational dynamics have never been experimentally characterized. Without structural data and with no single defined glycoform to model, empirical screening cannot enrich for binding against an invisible target, and machine learning cannot learn from zero samples. At the root of this impasse lies a universal deficit of causal transparency [20]: current methods share a common prerequisite, the availability of training data, and when structural data are absent, they cannot model, optimize, or uncover functional mechanisms [21, 22], let alone access a deeper physical possibility: that an intrinsically disordered glycan shield might, within a suitably engineered interface, be remodeled into an ordered conformation and functionally inverted into a binding anchor.

Overcoming this limitation demands a framework that does not require complete structures, but instead leverages causal inference and physics-based simulation to explore functional mechanisms from limited prior knowledge. In this paper we introduce the Antibody Evolution Nexus with Causal-Driven Simulation (AENCS), a white-box framework integrating molecular simulation, causal inference, and experimental validation within a unified workflow (Fig. 1a). Applying AENCS to restore S309 efficacy against JN.1 [8, 10], and guided by little more than the knowledge of the N354 glycosylation site, we identified ACC01, an optimized antibody exhibiting∼24-fold improved neutralization potency. Meanwhile, ACC01 spontaneously stabilized the otherwise disordered N354 glycan [7], enabling its first cryo-EM atomic-resolution definition and mechanistic elucidation. Quantitative causal analysis and structural dissection further revealed how this glycan shield is functionally inverted into an auxiliary binding anchor (Fig. 1b) through a multi-layered interaction network. This mechanistic understanding proved predictive: given the conservation of N354 glycosylation since its acquisition, ACC01 should retain its activity against emerging variants. Experimental validation confirmed this prediction, with ACC01 maintaining potent neutralization against the latest variant NB.1.8.1 (Fig. 1c). This trajectory from affinity optimization to mechanistic discovery to cross-variant protection demonstrates that causal-driven antibody engineering elevates mechanistic insight into functional utility, transforming antibody development from empirical screening into a knowledge-driven pursuit, and providing a viable paradigm for exploring the long-vacant frontier of antibody design targeting glycosylated epitopes.

**Figure 1.**
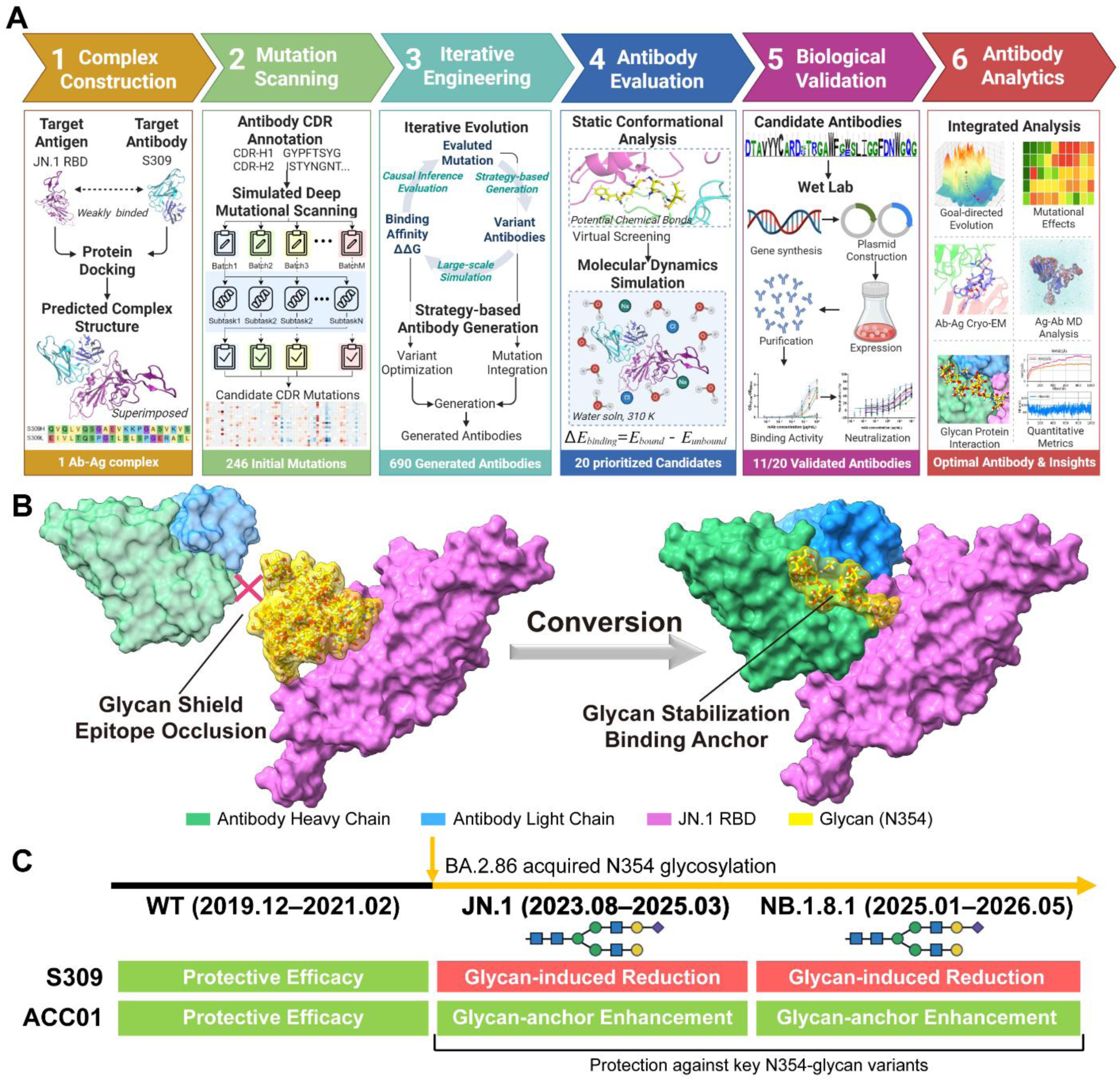
AENCS computational-experimental framework and glycan shield-to-anchor conversion mediates broad-spectrum neutralization by ACC01. **(a)** Schematic workflow of the AENCS framework for antibody optimization. The six integrated stages span computational design (Stages 1–4), experimental validation (Stage 5), and mechanistic analysis (Stage 6). Causal inference guides iterative evolution of the pre-experimental pipeline. **(b)** Structural depiction of the N354 glycan functional conversion. Left: In its unbound state, the N354 glycan on the JN.1 RBD (orchid) adopts multiple disordered conformations that form a steric shield, preventing antibody access. Right: In the ACC01/JN.1 RBD complex, the glycan is resolved in a single ordered conformation, serving as an auxiliary binding anchor. The arrow denotes the shield-to-anchor conversion. **(c)** Broad-spectrum neutralization profile of S309 and ACC01. The SARS-CoV-2 variants are ordered chronologically along the horizontal axis. S309 neutralizes only WT, losing activity against N354-bearing variants. ACC01 potently neutralizes all variants tested. The acquisition of the N354 glycosylation site is marked; variants carrying N354 are highlighted with glycan symbols. Abbreviations: RBD, receptor binding domain; CDR, complementarity-determining region; ΔΔG, change in Gibbs binding free energy; Ab–Ag, antibody–antigen; MD, molecular dynamics.

## Results

### AENCS enables effective antibody optimization

The AENCS framework integrates computational design, experimental validation, and mechanistic analysis within a unified, causality-driven workflow (Fig. 1a). The primary distinction lies in causal transparency, enabling the framework not only to generate optimized sequences but also to dissect the mechanistic basis of their enhanced function. The computational engine (Stages 1–4) initiates with template-guided modeling of the S309/JN.1-RBD complex (Stage 1), followed by parallel-accelerated deep mutational scanning across all complementarity-determining region positions [23] to identify affinity-enhancing single substitutions (Stage 2). Promising mutations seed several rounds of iterative evolution (Stage 3), wherein designed antibodies are generated, their binding energy evaluated via protein simulation, and causal filtering prioritizes genuinely causative mutations while disentangling them from spuriously correlated residues [24]. In the final generation phase, a slight hydrophobic perturbation was applied at heavy-chain position 108, which is spatially proximal to the N354 glycosylation site, yielding approximately 690 unique antibodies. Top-ranking antibodies subsequently undergo conformational filtering and molecular dynamics (MD) simulations to assess interface stability (Stage 4), yielding 20 candidates for experimental synthesis (Extended Data Table 1).

These 20 designed antibodies, together with S309, were subjected to comprehensive in vitro characterization (Stage 5; Fig. 2). ELISA binding assays against the extracellular domain of spike protein (S_ECD_) of SARS-CoV-2 wild-type (WT) and JN.1 demonstrated that all 21 antibodies cross-reacted with both antigens. It is noteworthy that 11 of the 20 designed antibodies exhibited enhanced binding to JN.1 S_ECD_ relative to S309, with greater than 5-fold improvements in half-maximal effective concentration (EC_50_) (Fig. 2a). Pseudovirus neutralization assays revealed that 8 of these 11 binding-enhanced designs displayed improved neutralization against JN.1, with all half-maximal inhibitory concentration (IC_50_) values below 20 μg/mL. ACC01 showed the most pronounced enhancement, achieving ∼24-fold improvement in IC_50_ (Fig. 2b). Thermal stability analysis showed that all 21 antibodies maintained melting temperatures above 60 ℃, with ACC01 and ACC02 exceeding 75 ℃, (Fig. 2c). ACC01, ACC02, and ACC04 were selected for authentic virus neutralization assays. All three antibodies potently neutralized both WT and JN.1 viruses, with ACC01 achieving the lowest IC_50_ (0.009 μg/mL against JN.1) (Fig. 2d). Surface plasmon resonance (SPR) measurements showed that all K_D_ values were below 10^−8^ M. For JN.1 S_ECD_, ACC01 exhibited at least a 2-fold higher binding response and over 3-fold improved K_D_ relative to S309 (Fig. 2e). Across all assays, ACC01 consistently ranked as the top performer.

**Figure 2.**
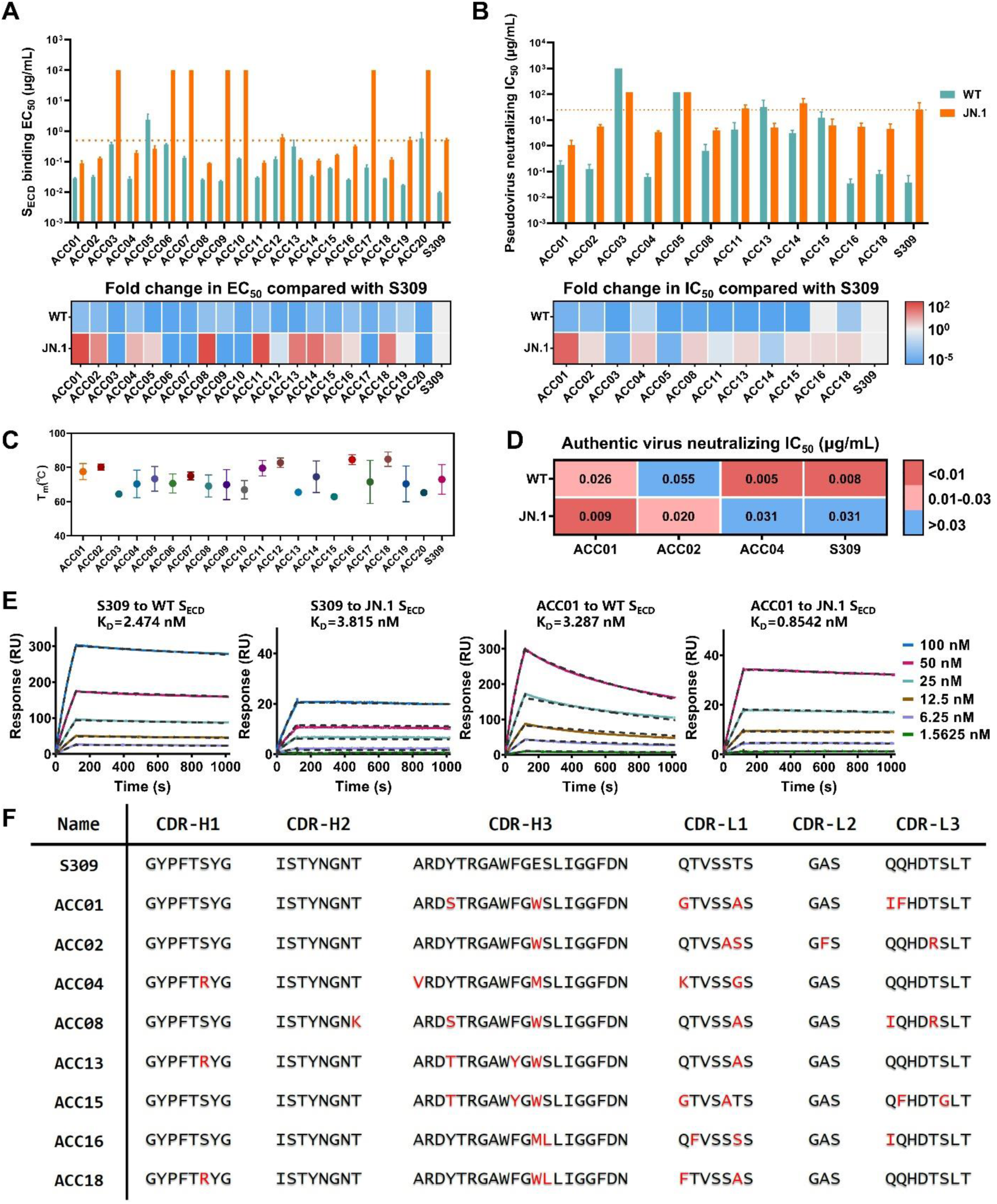
Functional characterization of AENCS-designed antibodies. (a) ELISA binding EC_50_ values of 20 designed antibodies and S309 to the spike extracellular domain (S_ECD_) of SARS-CoV-2 WT and JN.1 variants. Eleven antibodies exhibited enhanced JN.1 binding relative to S309, with fold changes indicated in the heatmap. All data are represented as means ± standard deviation (SD) of three replicates from one representative experiment. (b) Pseudovirus neutralization IC_50_ values of the 11 binding-enhanced antibodies and S309 against pseudotyped WT and JN.1 viruses, with fold changes relative to S309 indicated in heatmap. All data are represented as means ± SD of three replicates from one representative experiment. (c) Thermal stability analysis. Melting temperatures (T_m_) of the antibodies measured by the UNcle platform are shown. (d) Authentic virus neutralization IC_50_ values of ACC01, ACC02, ACC04 and S309 against WT and JN.1 viruses. Representative data from three independent experiments are shown. (e) Binding affinity of ACC01 and S309 to SARS-CoV-2 (WT) and JN.1 S_ECD_. Experimental responses (colored lines) and fitting curves using a 1:1 binding model (dotted lines) are shown. K_D_, equilibrium dissociation constant. (f) Sequence alignment of S309 and the 8 neutralization-enhanced antibodies including ACC01. AENCS-designed mutated residues are highlighted in red.

### Goal-directed efficiency and causal quantification

To evaluate goal-directed search capability, we reconstructed AENCS’s evolutionary trajectory and compared it against random mutagenesis. Across three rounds of increasing mutational complexity, AENCS designs converged efficiently toward the low-energy basin containing ACC01 (Extended Data Fig. 1). This convergence demonstrates that causal filtering, rather than exhaustive sampling, directs the search toward optimal solutions. The entire pre-experimental design completed within 28 hours, enabled by parallel acceleration (Extended Data Fig. 2).

AENCS also provides quantitative causal attribution for individual mutations. Using experimental binding data (EC_50_ fold change relative to S309; Extended Data Table 2), we estimated the Average Treatment Effect (ATE) for each mutation (Extended Data Table 3). All six ACC01 mutations exhibited positive ATEs, with heavy-chain E108W displaying the largest effect (ATE = 2.53), followed by others showing positive contributions (0.41–2.18). Notably, several mutations with favorable single-point predictions were absent from the optimal combination, confirming that functional enhancement arises from synergistic epistasis, precisely the rationale underlying AENCS’s causal filtering strategy.

### Enhanced direct antibody–antigen interactions through optimized interfacial networks

To elucidate the mechanistic basis for the enhanced function of ACC01, we determined the cryo-EM structures of S309 and ACC01 in complex with the JN.1 RBD respectively. The overall binding pose remained unchanged, with ACC01 engaging the same core epitope (Fig. 3a) [9]. We therefore focused on the protein–protein interface, performing 10-ns MD simulations.

**Figure 3.**
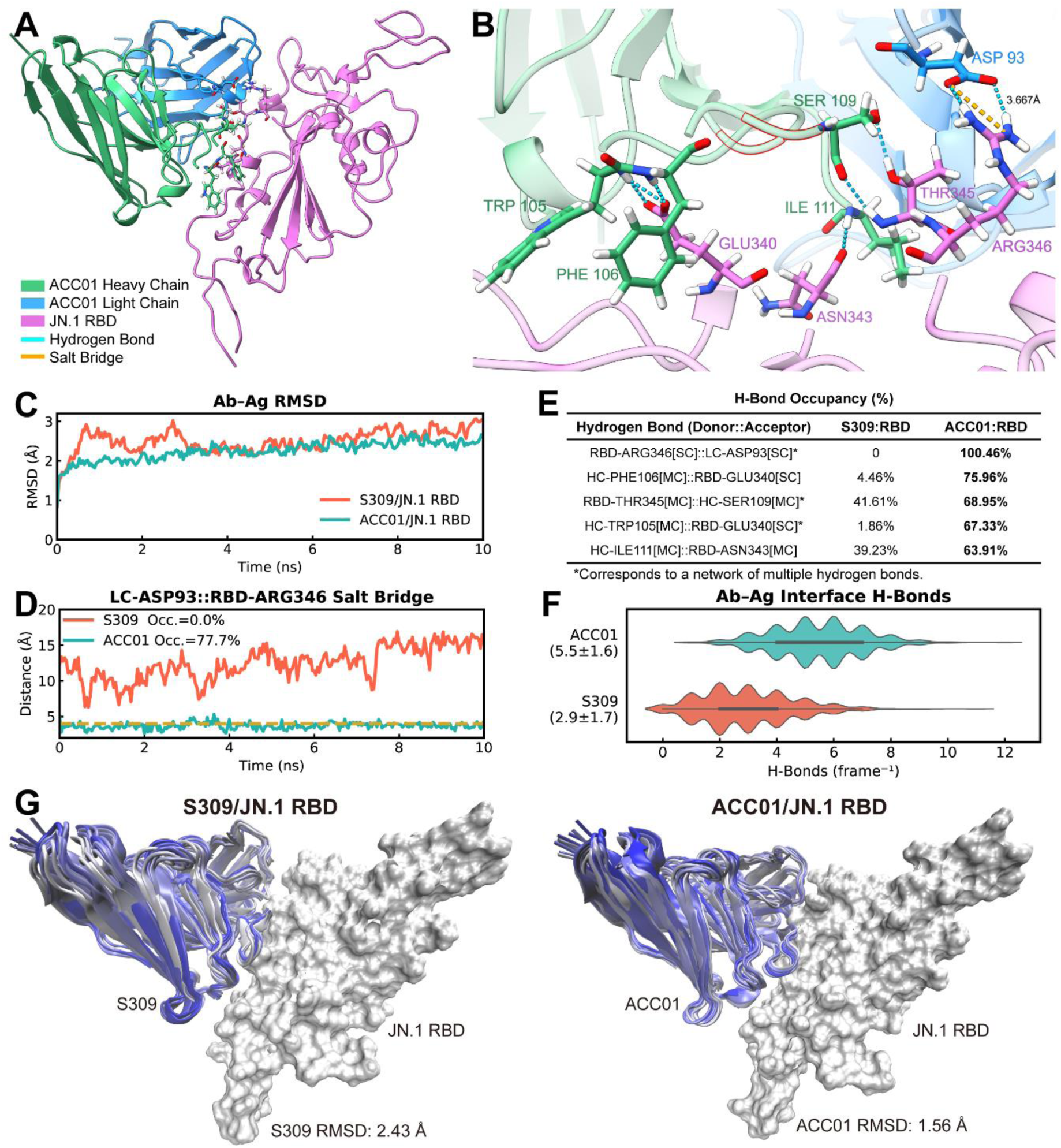
Pure protein interface: structural and dynamic basis for enhanced ACC01/JN.1 RBD complex stability. **(a)** Overall structure of the ACC01/JN.1 RBD complex. Antibody and RBD are shown in cartoon representations, with key interface residues highlighted as sticks. **(b)** High-resolution view of the interface showing key interactions. Hydrogen bonds (blue dashes; all distances <3.0 Å) and a salt bridge (orange dash) are indicated. For clarity, hydrogen bond distances are omitted due to the density of interactions. The snapshot corresponds to the 5-ns frame from the MD trajectory. Key residues are shown as sticks. Engineered residues are outlined in red. **(c)** RMSD of the ACC01/JN.1 RBD and S309/JN.1 RBD complexes over 10-ns MD simulations. ACC01 exhibits faster convergence and reduced fluctuations. Inset: potential and kinetic energy profiles (see Extended Data Fig. 3). **(d)** Salt bridge distance between light-chain Asp93 and RBD Arg346 over 10-ns MD simulations. The dashed line marks the 4.0 Å cutoff. Occupancy (Occ.) is indicated. **(e)** Occupancy comparison of key interfacial hydrogen bonds. Asterisks denote networks comprising multiple hydrogen bonds. **(f)** Total number of interfacial hydrogen bonds (mean ± s.d.) in ACC01 and S309 trajectories. **(g, h)** Conformational snapshots of S309/JN.1 RBD **(g)** and ACC01/JN.1 RBD **(h)** during the equilibrated phase of MD (1–10 ns). Frames were sampled every 1 ns and colored by time (silver → blue). Antibody RMSD values over this interval (reference: 1 ns) confirm the greater rigidity of ACC01. Abbreviations: RMSD, root-mean-square deviation; LC, light chain; HC, heavy chain; MC, main chain; SC, side chain.

ACC01 formed a markedly more stable complex with the JN.1 RBD than S309. All-atom RMSD revealed that the ACC01 complex converged more rapidly and exhibited greater overall stability, accompanied by a consistently lower potential energy (Fig. 3c; Extended Data Fig. 3). At the heart of this stabilization was a strengthened interfacial interaction network. A critical salt bridge between light-chain Asp93 and RBD Arg346, absent in S309, became firmly established in ACC01, with 77.7% occupancy (Fig. 3d) [25]. Concurrently, multiple hydrogen bonds anchoring the epitope were significantly reinforced (Fig. 3b). Hydrogen bonds involving RBD Glu340, Asn343, and Thr345, including those mediated by heavy-chain Trp105, Phe106, Ser109, and Ile111, exhibited occupancies ranging from ∼60% to ∼100%, whereas S309 occupancies remained below ∼40%. Notably, the engineered heavy-chain substitution E108W resided at the center of this interaction network, adjacent to Ser109 and in close contact with other residues, while Y100S lay nearby, positioning both mutations to potentiate the hydrogen bond network. Quantification of interface hydrogen bonds corroborated these findings: ACC01 averaged 5.5 hydrogen bonds, compared with 2.9 for S309 (Fig. 3f), representing a ∼90% increase.

Dynamic conformational sampling of the equilibrated trajectory further underscored the enhanced complementarity (Fig. 3g). Whereas S309 exhibited appreciable interfacial fluctuation and conformational drift (antibody RMSD = 2.43 Å), ACC01 remained tightly apposed to the RBD with minimal displacement (antibody RMSD = 1.56 Å), reflecting markedly reduced conformational heterogeneity. Thus, ACC01 achieved a more rigid, shape-complementary interface.

Collectively, ACC01 achieved enhanced binding by directly reinforcing interactions with the epitope, thereby counteracting the steric occlusion imposed by the N354 glycan. Consistent with ATE analysis, heavy-chain E108W is the principal driver of this interfacial remodeling. This augmented protein–protein interface provides the requisite physical foundation for stabilization of the N354 glycan.

### Structural discovery of the stabilized N354 glycan

Cryo-EM analysis of the ACC01/JN.1 RBD complex revealed a prominent structural feature. A continuous density map, morphologically characteristic of a carbohydrate chain, was apparent extending outward from the JN.1 RBD surface in the ACC01 complex, a density that, upon inspection, originated precisely from residue Asn354 (Fig. 4a and 4c, Extended Data Fig. 4). This is the N354-linked glycan, a glycosylation site acquired in BA.2.86 and retained in JN.1 [7], whose steric occlusion of key epitopes has been extensively documented [3]; however, its complete atomic structure has remained unresolved due to its inherent conformational disorder [2, 26]. In the S309 complex, by contrast, the glycan was largely invisible beyond faint, fragmented density at its base, reflecting the dynamic flexibility that has precluded structural characterization (Fig. 4a). Mass spectrometry identified a heterogeneous population of N-glycans at N354, with the predominant species corresponding to HexNAc₄Hex₅NeuAc₁ (Fig. 4b; Extended Data Fig. 4). Through orthogonal cryo-EM and mass spectrometric validation, we thus resolved the ordered core structure of the N354 glycan. The cryo-EM map accommodates the top three glycoforms, collectively accounting for ∼60% of the population (Extended Data Fig. 4):

Galβ1–4GlcNAcβ1–2Manα1–3(Manα1–6)Manβ1–4GlcNAcβ1–4GlcNAcβ1–N-Asn354

**Figure 4.**
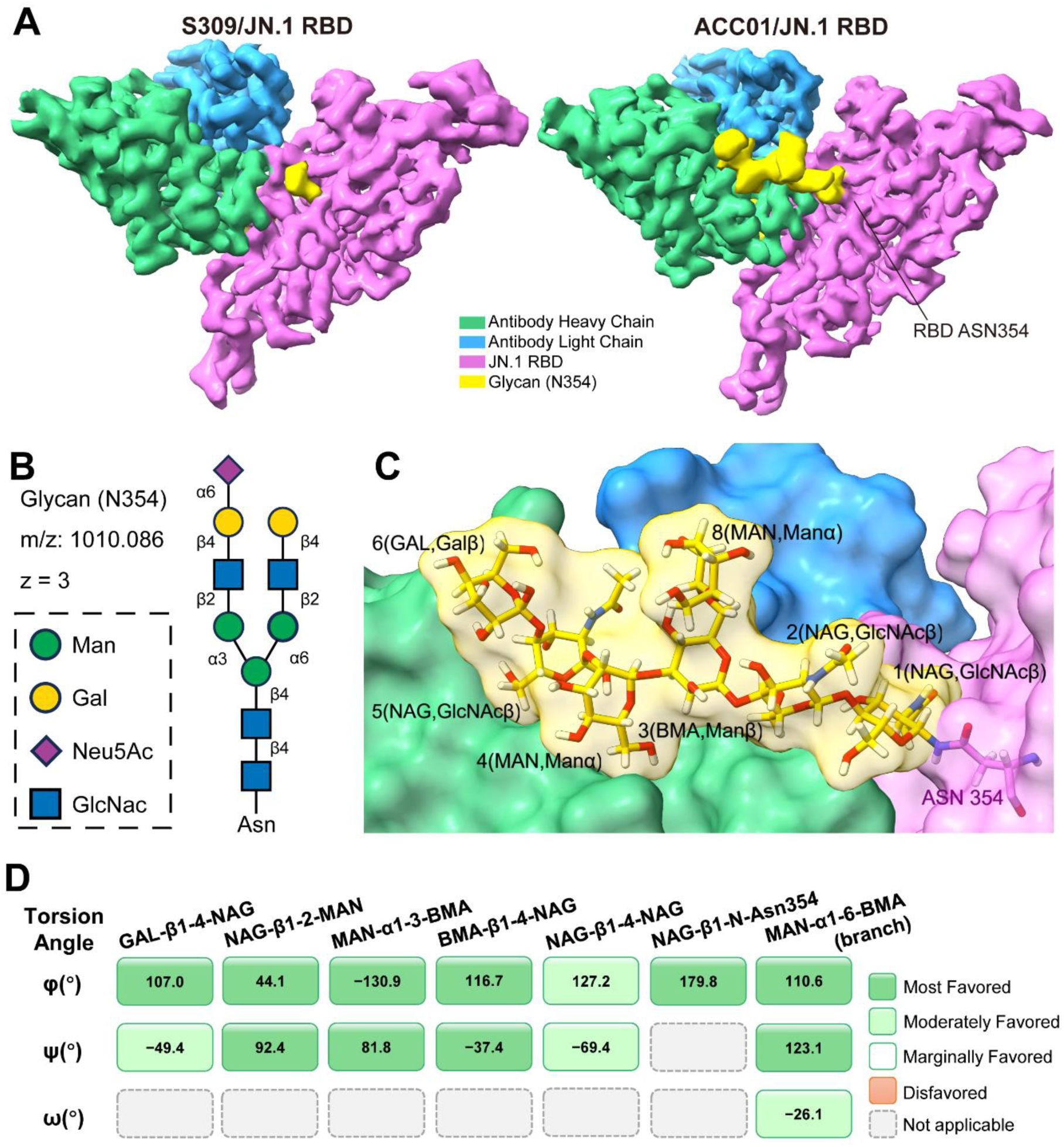
Cryo-EM and mass spectrometry reveal the structure and stabilization of the N354 glycan. **(a)** Cryo-EM densities of the N354 glycan in the S309/JN.1 RBD (left) and ACC01/JN.1 RBD (right) complexes. Only the ACC01 complex yields continuous, well-resolved density enabling atomic modeling of the complete glycan moiety. **(b)** Mass spectrometry identification of the N354-linked glycan. The spectrum shows the N-glycan composition attached to peptide FASVYAWNR (precursor *m/z* 1010.086, z=3). The schematic representation of the glycan is shown. **(c)** Close-up view of the N-linked glycan at Asn354 of JN.1 RBD. The glycan is shown as sticks, with full sequence Galβ1–4GlcNAcβ1–2Manα1–3(Manα1–6)Manβ1–4GlcNAcβ1–4GlcNAcβ1–N-Asn354. Residue labels in the figure indicate the residue number and name. The glycan is stabilized in a relatively open region on the side of the antibody heavy chain, away from the core epitope, thus avoiding steric occlusion of the antigen-binding interface. **(d)** Glycosidic torsion angle validation for the N354 glycan. All φ, ψ, and ω torsion angles fall within energetically favorable ranges for unstrained N-glycans (see Extended Data Table 4 for full details). Color coding: dark green = Most Favored (deviation ≤0.33 from range center); light green = Moderately Favored (0.33–0.67). No disfavored angles were observed. Abbreviations: GlcNAc = N-acetylglucosamine (NAG), Man = mannose (MAN/BMA), Gal = galactose (GAL), Asn = asparagine.

The glycan is accommodated within a pocket formed jointly by ACC01 and RBD (Fig. 4c). Critically, this binding site is spatially segregated from the core S309 epitope, thereby avoiding steric interference with antigen engagement. Validation of glycosidic torsion angles confirmed that all φ, ψ, and ω values reside within the low-energy conformational landscape of unstrained N-glycans (Fig. 4d; Extended Data Table 4) [27].

Collectively, guided by little more than the knowledge of the N354 glycosylation site, ACC01 not only augmented the direct protein–protein interface but also spontaneously stabilized the otherwise dynamically disordered N354 glycan. This ordering countered a key immune-evasion element and unveiled a new dimension of the binding interface.

### Molecular mechanism and functional contribution of glycan stabilization

The stabilization of the N354 glycan by ACC01 arises from a multi-layered interaction network (Fig. 5).

**Figure 5.**
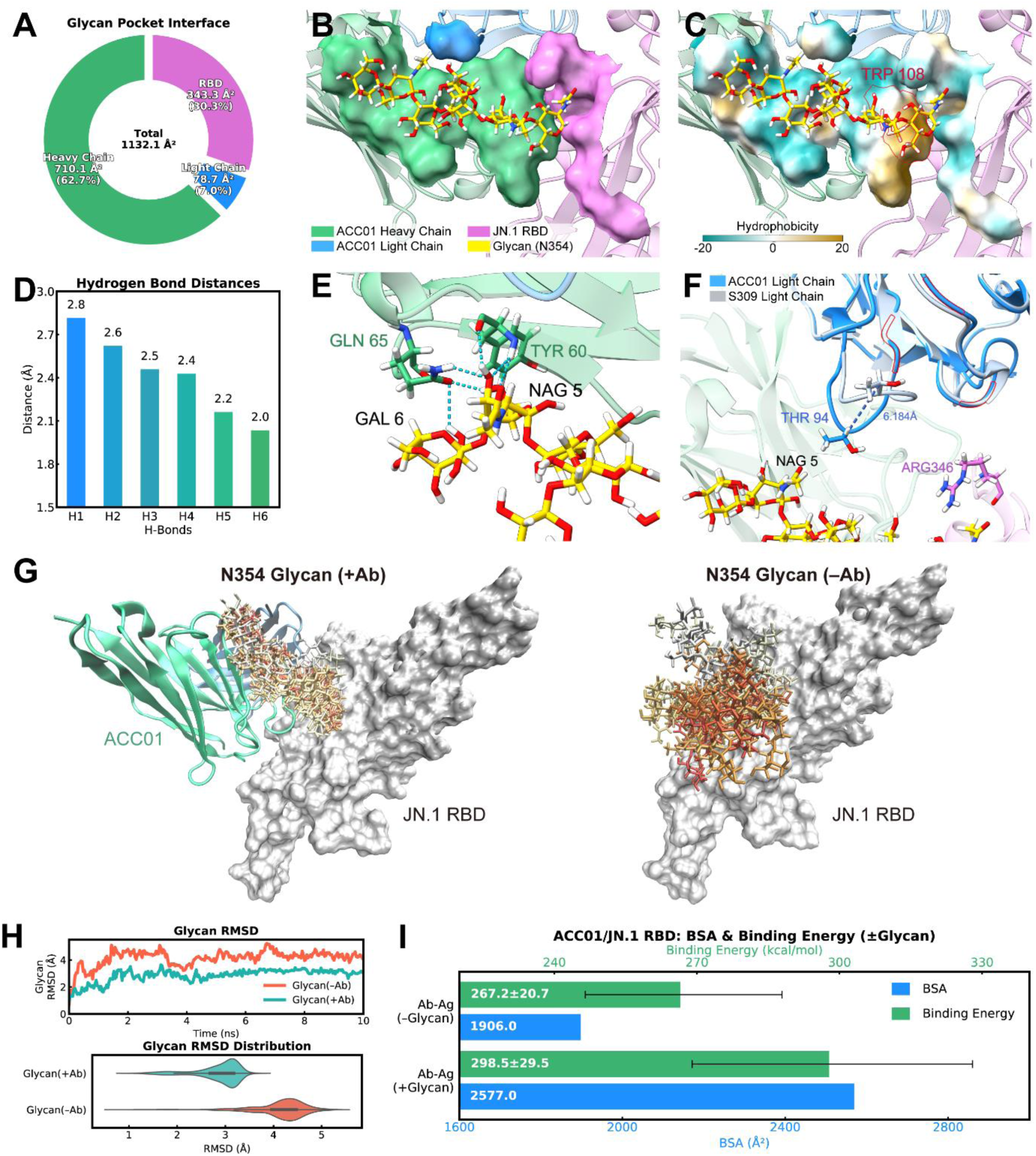
Mechanism of glycan stabilization and its energetic contribution to complex formation. **(a)** Contact area contributed by each component to the 5-Å glycan-interacting pocket. Percentages are indicated. **(b)** Visualization of the 5-Å pocket, colored by chain (heavy chain, green; light chain, blue; RBD, orchid). The antibody heavy chain provides the majority of van der Waals contacts. **(c)** Hydrophobicity mapping of the pocket. E108W (red) abuts the glycan core and accounts for ∼50% of the local hydrophobic surface. **(d)** Distances of six hydrogen bonds between glycan termini (GAL6, NAG5) and antibody heavy chain Gln65/Tyr60. All distances are <3.0 Å. **(e)** Atomic detail of the hydrogen bonds (blue dashes) shown in **(d)**. **(f)** Conformational change in light chain induced by distal mutations. Alignment of ACC01 (colored) with S309 (gray) shows reorientation of the Asp93–Ser95 turn. Thr94 shifts ∼6 Å toward the glycan. Mutated residues (T32A, Q90I, Q91F, Q27G) are in red. **(g)** Conformational dynamics of N354 glycan with (left) and without (right) ACC01. Snapshots sampled at 1-ns intervals are superimposed and colored by simulation time (white → red). **(h)** RMSD of the glycan with and without antibody. Means ± standard deviation is indicated; inset shows violin plot of RMSD distributions over simulation time. **(i)** Contribution of the N354 glycan to complex formation. Top: Buried surface area (BSA) increases by 671 Å² (35%). Bottom: Binding energy calculated over 10-ns MD. Inclusion of the glycan substantially contributes to binding energy. Abbreviations: RMSD, root-mean-square deviation; BSA, buried surface area.

The glycan is accommodated within a 1,132 Å² pocket shaped jointly by the heavy chain (62.7%), light chain, and RBD (Fig. 5a, b). While the pocket is largely hydrophilic, a conspicuous hydrophobic patch dominates the glycan-proximal region. Here, the engineered heavy-chain substitution E108W (ATE = 2.53), the single most decisive modification for glycan stabilization, directly abuts the glycan core, accounting for ∼50% of the local hydrophobic surface, and forms, together with Phe106, a continuous aromatic ridge that cradles the proximal saccharides, consistent with a dominant hydrophobic contribution rather than an electrostatic effect (Fig. 5c, Extended Data Fig. 5) [26].

In concert with this hydrophobic anchor, a latent polar network is awakened. Two heavy-chain residues, Gln65 and Tyr60, engage the Galβ1–4GlcNAc through six hydrogen bonds, with all distances below 3.0 Å (Fig. 5d, e). Absent in the wild-type context, these interactions are unmasked by the global interface remodeling achieved by ACC01, providing a second, polar tether that locks the glycan terminus.

Meanwhile, distal light-chain mutations provide additional refinement, inducing a concerted main chain displacement of the Asp93–Ser95 turn, shifting Thr94 by approximately 6 Å (Fig. 5f). This repositioning brings Thr94 into the glycan-contacting pocket while simultaneously reinforcing a salt bridge between Asp93 and Arg346 and hydrogen-bond network (Fig. 3). The light-chain ATE values (0.41–2.18) are thus rationalized.

To assess the dynamic consequences of this interaction network, we performed 10-ns MD simulations of the glycosylated RBD in the presence and absence of ACC01. Without antibody, the N354 glycan sampled a broad, diffuse conformational ensemble (RMSD = 4.1 Å), its terminus sweeping through a volume that readily occluded the core epitope (Fig. 5g, h). In contrast, antibody binding collapsed this heterogeneity: the glycan adopted a tightly restricted, well-ordered state (RMSD = 2.8 Å) and maintained a consistent orientation that avoided occlusion of the core epitope.

Critically, the stabilized glycan is not merely a passive occupant; it actively fortifies the interface. Inclusion of the N354 glycan expanded the buried surface areas (BSA) by 35% (671 Å²) and augmented the computed binding energy by 31.3 kcal/mol (Fig. 5i). This substantial energetic gain, which far exceeds typical non-specific glycan contributions [26], reflects the multi-dimensional mechanism described above: hydrophobic anchoring at the glycan base (E108W/Phe106) and hydrogen-bond tethering at its terminus (Gln65/Tyr60) provide the specific, high-affinity contacts, while extensive van der Waals encapsulation by the heavy chain (accounting for 92% of all atomic contacts; Extended Data Table 5) further stabilizes the complex. The N354 glycan, originally a steric shield, is thus functionally inverted into an auxiliary binding anchor [28]. This conversion, together with the reinforced protein–protein interface, establishes the mechanistic foundation for ACC01’s enhanced neutralization potency.

### ACC01 retains potent activity against the latest variant NB.1.8.1

The functional conversion of the N354 glycan into a binding anchor prompted us to examine its conservation across the BA.2.86 lineage. The N354 glycosylation site has been stably inherited by all subsequent variants, including the recently dominant NB.1.8.1, which has circulated persistently since early 2025 (Fig. 6a) [29, 30]. Structural mapping revealed that the mutations accumulated in NB.1.8.1 do not encroach upon the glycan-binding pocket or the core epitope, while sequence alignment confirmed the conservation of both the epitope and the N354 glycosylation site (Fig. 6b–c). Together, these findings predict that ACC01 should retain activity against NB.1.8.1.

**Figure 6.**
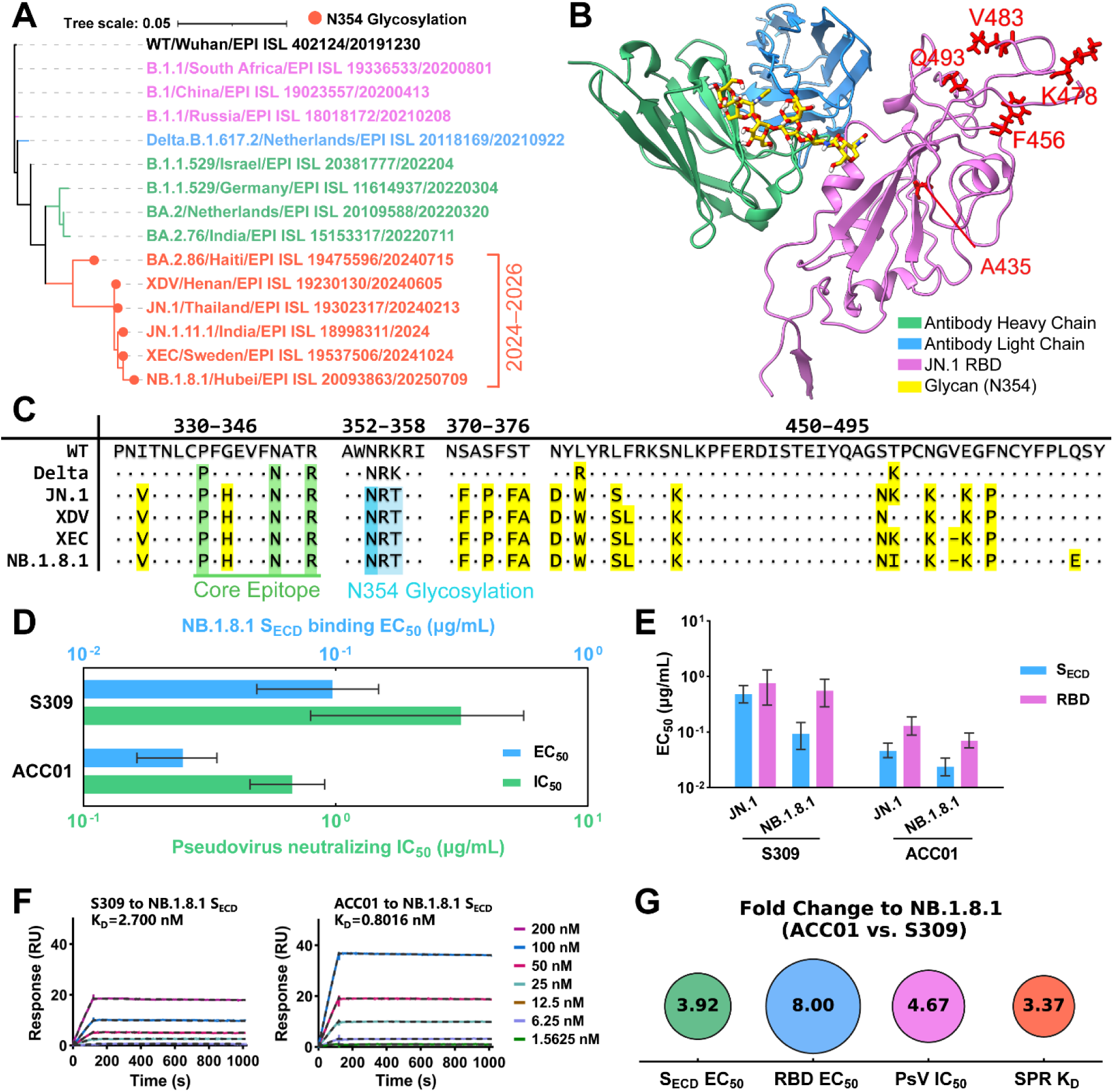
Evolutionary conservation and structural basis for ACC01 activity against circulating NB.1.8.1 variant. **(a)** Phylogenetic relationships of major SARS-CoV-2 variants, highlighting the emergence of the N354 glycosylation site in BA.2.86 and its conservation in subsequent variants. Filled circles denote N354-bearing variants. **(b)** Structural mapping of NB.1.8.1 RBD mutations (red sticks) onto the ACC01/JN.1 RBD complex. NB.1.8.1 mutations do not encroach upon the glycan-binding pocket or the core epitope, and are unlikely to affect glycan engagement significantly. **(c)** Multiple sequence alignment of key RBD regions. Both the core epitope and the N354 glycosylation site are generally conserved among the N354-bearing variants. **(d)** Binding EC_50_ values of S309 and ACC01 to NB.1.8.1 S_ECD_ and neutralization IC_50_ values against NB.1.8.1 pseudotyped variant. **(e)** Binding EC_50_ values of S309 and ACC01 against S_ECD_ and RBD of JN.1 and NB.1.8.1. **(f)** Binding affinity of ACC01 and S309 to NB.1.8.1 S_ECD_ measured by SPR assay. Experimental responses (colored lines) and fitting curves using a 1:1 binding model (black dotted lines) are shown. K_D_, equilibrium dissociation constant. **(g)** Fold change of ACC01 relative to S309 in the four measured parameters: S_ECD_ binding EC_50_, RBD binding EC_50_, pseudovirus neutralization IC_50_, and SPR K_D_ against NB.1.8.1. Abbreviations: PsV, pseudovirus; S_ECD_, spike extracellular domain; RBD, receptor-binding domain.

We therefore measured the binding and neutralization activities of ACC01 and S309 against NB.1.8.1. ACC01 exhibited substantially improved binding to both S_ECD_ and RBD, with EC_50_ enhancements of 3.9-fold and 8.0-fold, respectively, and maintained a 4.7-fold improvement in pseudovirus neutralization (Fig. 6d–g). These results confirm that this mechanistic understanding is predictive: by converting the conserved N354 glycan into a binding anchor, ACC01 extends its protective activity to the most recent dominant variant, surpassing its parental antibody in both potency and durability of protection.

Collectively, ACC01 not only resists glycan-mediated occlusion through a reinforced protein–protein interface (Fig. 3), but also captures and stabilizes the N354 glycan (Fig. 4), co-opting it as a binding enhancer (Fig. 5). Extending this mechanistic understanding beyond the original target, ACC01 converts the conserved N354 glycan into a durable binding anchor, achieving potent neutralization against the latest dominant variant NB.1.8.1 (Fig. 6). This trajectory from affinity optimization to mechanistic discovery to cross-variant protection demonstrates that causal-driven antibody engineering can translate mechanistic insight into functional utility, providing a viable strategy for converting conserved glycans into durable protective anchors against evolving variants.

## Discussion

This research applied AENCS to optimize S309 against the immune-evasive JN.1 variant, yielding ACC01 with ∼24-fold improved neutralization. Although AENCS operated with only the knowledge that N354 is a glycosylation site, the stabilization, structural resolution, and functional conversion of this glycan into a binding anchor rest on three sequential mechanistic pillars. First, the optimization efficiently identified ACC01 that established a reinforced protein–protein interface, providing the requisite physical foundation for resisting glycan-mediated steric occlusion (Fig. 3). Second, ATE quantification pinpointed heavy-chain E108W as the dominant driver, while atomic-level dissection elucidated how this hydrophobic anchor, together with an awakened polar network and distal conformational cooperativity, captures the disordered glycan and locks it into a defined conformation. Third, this captured glycan is functionally exploited, significantly contributing to BSA and binding energy, converting a rapidly evolving immune-evasion element into an auxiliary binding anchor (Fig. 4–5). This trajectory from resistance to capture to exploitation demonstrates that causal-driven antibody optimization can unlock latent structural features, achieving what remains inaccessible to both empirical screening and black-box prediction when structural data are absent.

This mechanistic understanding enabled a testable prediction: given the conservation of the N354 glycan across the BA.2.86 lineage and that later mutations do not significantly perturb the binding interface, ACC01 should retain activity against the most recent emerging variants. Experimental validation confirmed this prediction, with ACC01 maintaining potent neutralization against NB.1.8.1, the dominant variant circulating persistently for years since early 2025 (Fig. 6). Notably, this predictive capacity holds despite glycoform heterogeneity at the N354 site, underscoring that causal-driven frameworks can generate transferable knowledge from limited prior information even when the target structure is not uniquely defined. This capacity to generate and validate mechanistic predictions from very limited prior knowledge distinguishes causal-driven frameworks from conventional approaches: AENCS yields transferable mechanistic knowledge that remains actionable against viral evolution, even when the target structure has never been experimentally resolved. Causal transparency thus transforms antibody engineering from retrospective optimization into prospective discovery.

This work establishes that causal-driven antibody engineering can operate effectively in a domain where rational design has long been stalled: the targeting of glycosylated epitopes. The scarcity of glycan structural data, compounded by inherent glycoform heterogeneity, has rendered systematic antibody design against these targets virtually impossible. AENCS addresses this impasse not by requiring complete structural information, but by leveraging causal inference and physics-based simulation to explore and exploit functional mechanisms in this data-sparse context. The discovery that a steric glycan shield can be converted into a protective binding anchor, validated by durable neutralization of the latest dominant variant, demonstrates that glycosylated epitopes can be systematically targeted through causal-driven discovery. By embedding causal inference within a unified computational-experimental workflow, AENCS redefines antibody optimization as a knowledge-driven discipline, providing a viable paradigm for illuminating cryptic binding determinants such as glycosylated epitopes and translating them into therapeutic strategies against evolving pathogens.

## Data and code availability

The atomic coordinates and cryo-EM density maps have been deposited in the RCSB PDB and EMDB. All other data and code supporting this study will be made publicly available at Zenodo upon publication. Detailed methods are provided in the Supplementary Information, which will be available in the final published version.

## Biosafety and Ethics Statement

This study did not involve human subjects, human biological samples, or animal experiments; thus, ethical approval was not required for the present work. All routine experimental procedures, including antibody expression and purification, enzyme-linked immunosorbent assay (ELISA), surface plasmon resonance (SPR) detection, and protein thermal stability analysis, were performed in accordance with standard laboratory operating specifications. Pseudotyped virus neutralization assays were carried out in a Biosafety Level 2 (BSL-2) laboratory following institutional approved guidelines. Authentic SARS-CoV-2 virus neutralization experiments were conducted in a Biosafety Level 3 (BSL-3) laboratory with official project authorization, and all operations complied with the institutional standard operating procedures for highly pathogenic respiratory viruses.

## Funding

This work was supported by the National Natural Science Foundation of China (82522044 and 62206309).

## Acknowledgments

The authors acknowledge the authors and laboratories that shared PDB structures and genomic sequences via RCSB and GISAID, on which this research is based.

## Author contributions

X.W., Conceptualization, Methodology, Writing - original draft; G.Z., Validation, Writing - original draft; M.H., Methodology, Writing - review & editing; J.H., Validation, Writing - original draft; C.S., Validation, Formal analysis; L.Z., Investigation, Formal analysis; Z.C., Resources; P.H., Investigation; W.W., Data curation; X.Z., Visualization; Y.D., Investigation; Y.Z., Investigation; P.L., Data curation; X.Z., Data curation; R.J., Writing - review & editing; H.W., Writing - review & editing; C.W., Writing - review & editing; X.L., Validation, Resources; L.Y., Formal analysis, Resources; Z.L., Formal analysis, Resources; H.R., Conceptualization, Supervision; J.X., Project administration, Supervision; X.C., Conceptualization, Funding acquisition, Writing - review & editing.

## Competing interests

The authors declare that a patent application related to the ACC01 antibody described in this work has been filed (application number: [202610799074.9]). All other authors declare no competing interests.

## Extended Data Tables and Figures

**Extended Data Table 1.** Candidates of designed antibodies selected for wet-lab validation alongside the original antibody (S309).

| Record ID | Heavy Chain Mutations | Light Chain Mutations |
| --- | --- | --- |
| ACC01 | Y100S, E108W | T32A, Q90I, Q91F, Q27G |
| ACC02 | E108W | S31A, T32S, T94R, A52F |
| ACC03 | R102I, Y100T, S31L, F114M, S109D | Q27K, Q90I, A52F, T32P |
| ACC04 | E108M, A97V, S31R | Q27K, T32G |
| ACC05 | F114M, F106Y, Y100T, E108W, S31R | T32A |
| ACC06 | Y100Q, Y27W, A97V, S31R | None |
| ACC07 | T58R, S109D, F106Y, S31N | S31A, Q90L, T28S |
| ACC08 | Y100S, T58K, E108W | T32A, T94R, Q90I |
| ACC09 | Y100S, S109W | A52Q, Q27S, Q91L |
| ACC10 | Y100N, S31L, F114M, S109D | Q27K, Q90L, T32P, A52M |
| ACC11 | Y100S, E108W, T58R, S31R | Q27G, T32A, Q91F, Q90I |
| ACC12 | T58R, S31R, E108K, A97I | A52M, T28A, T32S |
| ACC13 | Y100T, F106Y, S31R, E108W | T32A |
| ACC14 | Y100Q, T58R, E108W | T32N |
| ACC15 | Y100T, F106Y, E108W | S95G, S31A, Q91F, Q27G |
| ACC16 | E108M, S109L | T28F, Q90I, T32S |
| ACC17 | Y100N, S31L, F114M | Q27K, Q90I, Q91F, T32P, A52M, T94R |
| ACC18 | S31R, E108W, S109L | Q27F, T32A |
| ACC19 | None | T94K, Q90I, Q91M, T32A |
| ACC20 | Y100N, S31R, F106Y | Y100N, F106Y, T32P |
| S309 | None | None |

*Reference sequences:*
*S309 Variable Heavy Chain (VH):*
QVQLVQSGAEVKKPGASVKVSCKASGYPTSYGISWVRQAPGQGLEWMGWISTYNGNTNYAQKFQ GRVTMTTDTSTTTGYMELRRLRSDDTAVYYCARDYTRGAWFGESLIGGFDNWGQGTLVTVSS
*S309 Variable Light Chain (VL):*
EIVLTQSPGTLSPGERATLSCRASQTVSSTSLAWYQQKPGQAPRLLIYGASSRATGIPDRFSGSGSGT DFTLTISRLEPEDFAVYYCQHQHDTSLTFGGGTKVEIKR
Mutation format: [Original AA][Position][Mutated AA]. Residue numbering: IMGT scheme.

**Extended Data Figure 1.**
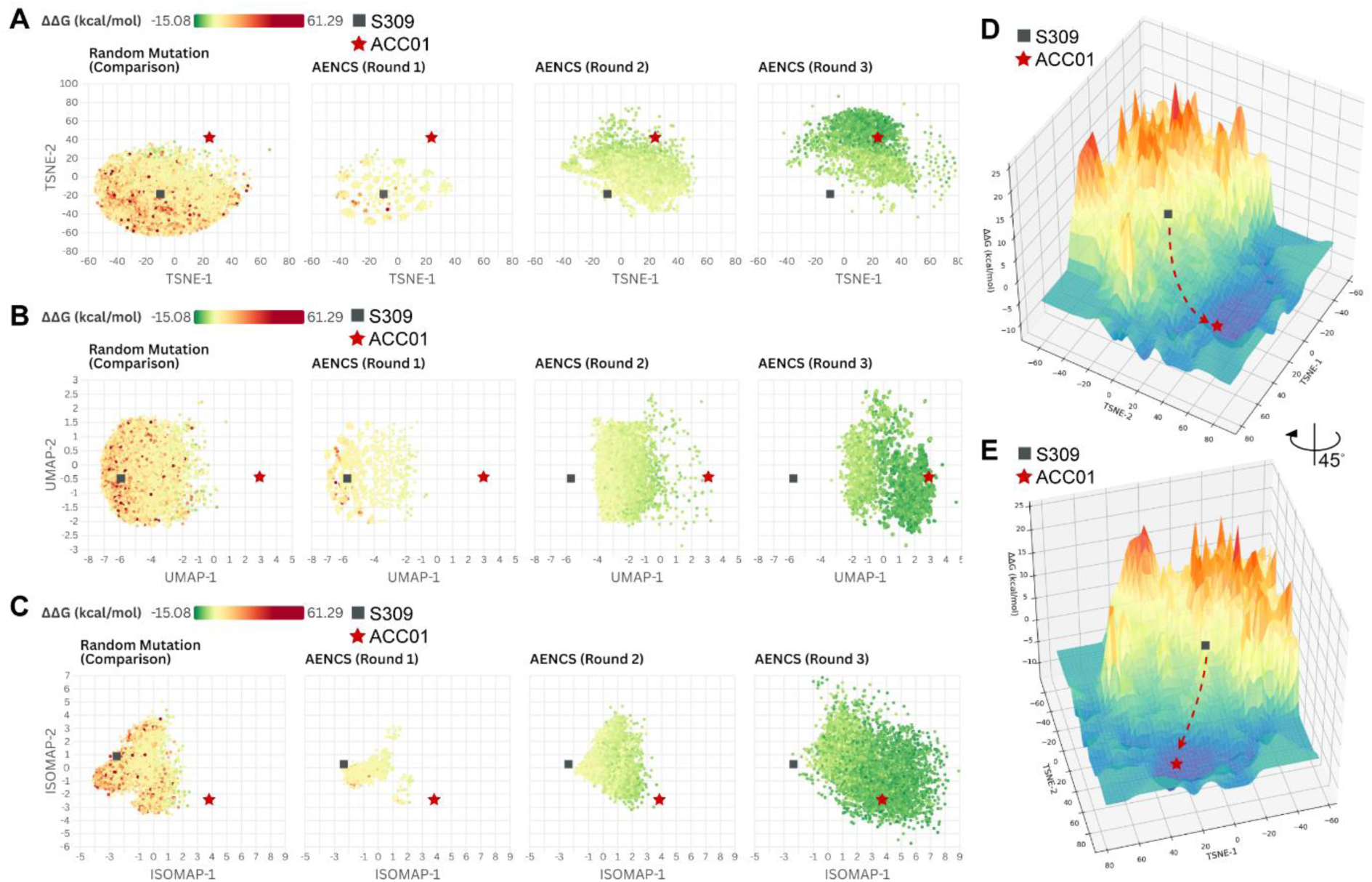
AENCS enables efficient goal-directed navigation to high-affinity antibodies. **(a–c)** Dimensionality reduction visualization of mutant antibody distributions across evolution strategies: random mutagenesis (10,000 antibodies), and iterative AENCS rounds (Round 1: 1,026 antibodies; Rounds 2–3: 5,000 antibodies per round). Antibodies are encoded by sequence features, with node colors indicating the antibody–antigen binding free energy change (ΔΔG, kcal/mol) calculated by FoldX; lower ΔΔG values correspond to stronger predicted affinity. Visualizations were generated via **(a)** t-SNE, **(b)** UMAP, and **(c)** IsoMap embedding. **(d, e)** Directional convergence of AENCS-guided antibody populations mapped onto the antibody sequence affinity landscape (t-SNE embedding), colored by binding ΔΔG. The S309 antibody is indicated by a red square (▪), and the optimized high-affinity antibody ACC01 is indicated by a red star (★).

**Extended Data Figure 2.**
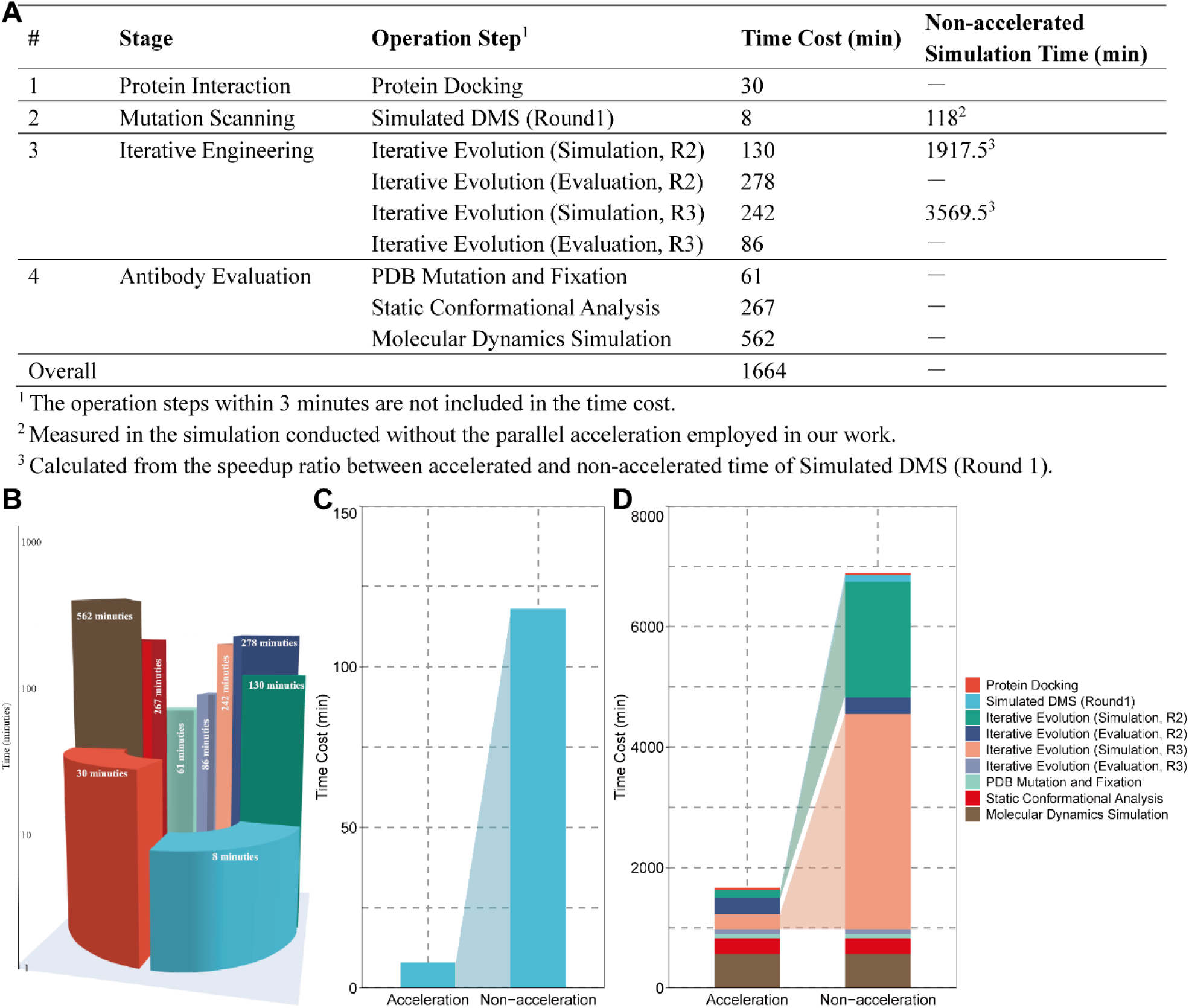
Computational time analysis of the AENCS *in silico* workflow (Stages 1–4). **(a)** Tabulated breakdown of time costs for individual operations. Key simulation steps (e.g., Simulated DMS, Iterative Rounds R2 & R3) are annotated with both the accelerated (parallel) duration and the estimated non-accelerated (serial) baseline duration. **(b)** Visualization of step-specific time costs, highlighting variations across stages. **(c)** Direct comparison of accelerated versus non-accelerated processing modes for the Simulated Deep Mutational Scanning (DMS) step, showing a 14.75-fold speedup. **(d)** Stacked bar plot summarizing cumulative time expenditure across all simulation steps, contrasting the total accelerated workflow duration (27.7 hours) against the projected non-accelerated total (114.8 hours) according to the 14.75-fold simulation speedup. Colors correspond to specific operation steps.

**Extended Data Table 2.** Experimental binding data for antibodies to JN.1 Spike. Fold change in EC_50_ relative to S309 for the 20 AENCS-designed antibodies. Antibodies with EC_50_ > 100 µg/mL were are reported as >100 and considered non-binding, and their fold change is reported as not applicable (NA). Antibody sequences are detailed in Extended Data Table 1. These binding data serve as the phenotypic input for the causal inference analysis in Extended Data Table 3.

| Record ID | EC<sub>50</sub> (ug/mL) | Fold change to S309 (EC<sub>50</sub>) |
| --- | --- | --- |
| ACC01 | 0.088436667 | 5.599110475 |
| ACC02 | 0.129233333 | 3.831570802 |
| ACC03 | >100 | NA |
| ACC04 | 0.189866667 | 2.607970506 |
| ACC05 | 0.260966667 | 1.897432622 |
| ACC06 | >100 | NA |
| ACC07 | >100 | NA |
| ACC08 | 0.088923333 | 5.568467219 |
| ACC09 | >100 | NA |
| ACC10 | >100 | NA |
| ACC11 | 0.09179 | 5.394560046 |
| ACC12 | 0.616533333 | 0.803146626 |
| ACC13 | 0.115566667 | 4.284684165 |
| ACC14 | 0.109016667 | 4.542118942 |
| ACC15 | 0.1635 | 3.028542304 |
| ACC16 | 0.3188 | 1.55322041 |
| ACC17 | >100 | NA |
| ACC18 | 0.116126667 | 4.264022045 |
| ACC19 | 0.5069 | 0.976852765 |
| ACC20 | >100 | NA |
| S309 | 0.495166667 | 1 |

**Extended Data Table 3.** Causal contribution of individual mutations to enhanced binding. Average treatment effect (ATE) of each mutation on fold change in binding affinity, estimated using Double Machine Learning. Because this analysis included only antibodies with measurable binding improvements, the ATE values reflect relative contributions within a functionally enhanced panel: positive values indicate stronger contributions to binding enhancement, while negative values indicate relatively weaker contributions rather than intrinsically deleterious effects. The mutations present in ACC01 are highlighted in bold. These results directly validate AENCS’s causal transparency: all mutations in ACC01 are genuine contributors to functional improvement, and their ATE ranking (heavy-chain E108W highest) aligns with their structural roles (Fig. 3–5). Abbreviations: ATE, average treatment effect; CI, confidence interval.

| **Mutation\*** | **ATE** | **90% CI** |
| --- | --- | --- |
| **EH108W** | 2.5253 | (1.3904, 3.6602) |
| **QL91F** | 2.1825 | (0.6366, 3.7284) |
| **YH100S** | 2.1333 | (1.3615, 2.9051) |
| **QL27G** | 1.6070 | (-0.7785, 3.9923) |
| TL32N | 1.3913 | (0.1271, 2.6555) |
| **QL90I** | 1.0461 | (-0.2288, 2.3210) |
| SH109L | 0.8575 | (-0.1617, 1.8766) |
| AH97V | 0.7872 | (0.7286, 0.8457) |
| AL52F | 0.6329 | (0.3849, 0.8809) |
| **TL32A** | 0.4141 | (-0.3031, 1.1312) |
| TL94R | 0.3874 | (-0.2610, 1.0359) |
| YH100Q | 0.3781 | (-1.4831, 2.2393) |
| SH31R | 0.2779 | (-0.5636, 1.1194) |
| QL27F | 0.0785 | (-0.7199, 0.8768) |
| TH58K | -0.1433 | (-1.3607, 1.0742) |
| TH58R | -0.2590 | (-1.5942, 1.0762) |
| FH106Y | -0.3829 | (-1.5112, 0.7455) |
| TL28F | -0.6056 | (-0.6322, -0.5789) |
| EH108M | -0.6768 | (-1.8950, 0.5414) |
| TL32G | -0.8600 | (-0.8624, -0.8576) |
| EH108K | -0.9005 | (-0.9021, -0.8989) |
| YH100T | -1.1163 | (-2.0772, -0.1554) |
| TL32S | -1.4911 | (-2.8438, -0.1384) |
| SL31A | -1.6075 | (-3.0741, -0.1409) |
| QL27K | -1.6523 | (-1.6546, -1.6500) |
| QL91M | -1.7315 | (-1.7326, -1.7304) |
| AH97I | -1.7324 | (-1.7335, -1.7314) |
| TL28A | -2.1609 | (-2.1627, -2.1590) |
| FH114M | -2.3587 | (-3.3128, -1.4046) |
| SL95G | -2.5411 | (-4.6414, -0.4408) |
| AL52M | -2.7426 | (-2.7437, -2.7416) |
| TL94K | -3.2834 | (-3.2857, -3.2811) |

\* Mutations are denoted as [Original residue][Chain][Position][Substituted residue]. For example, YH100S indicates that tyrosine (Y) at position 100 of H-chain is replaced by serine (S).

**Extended Data Figure 3.**
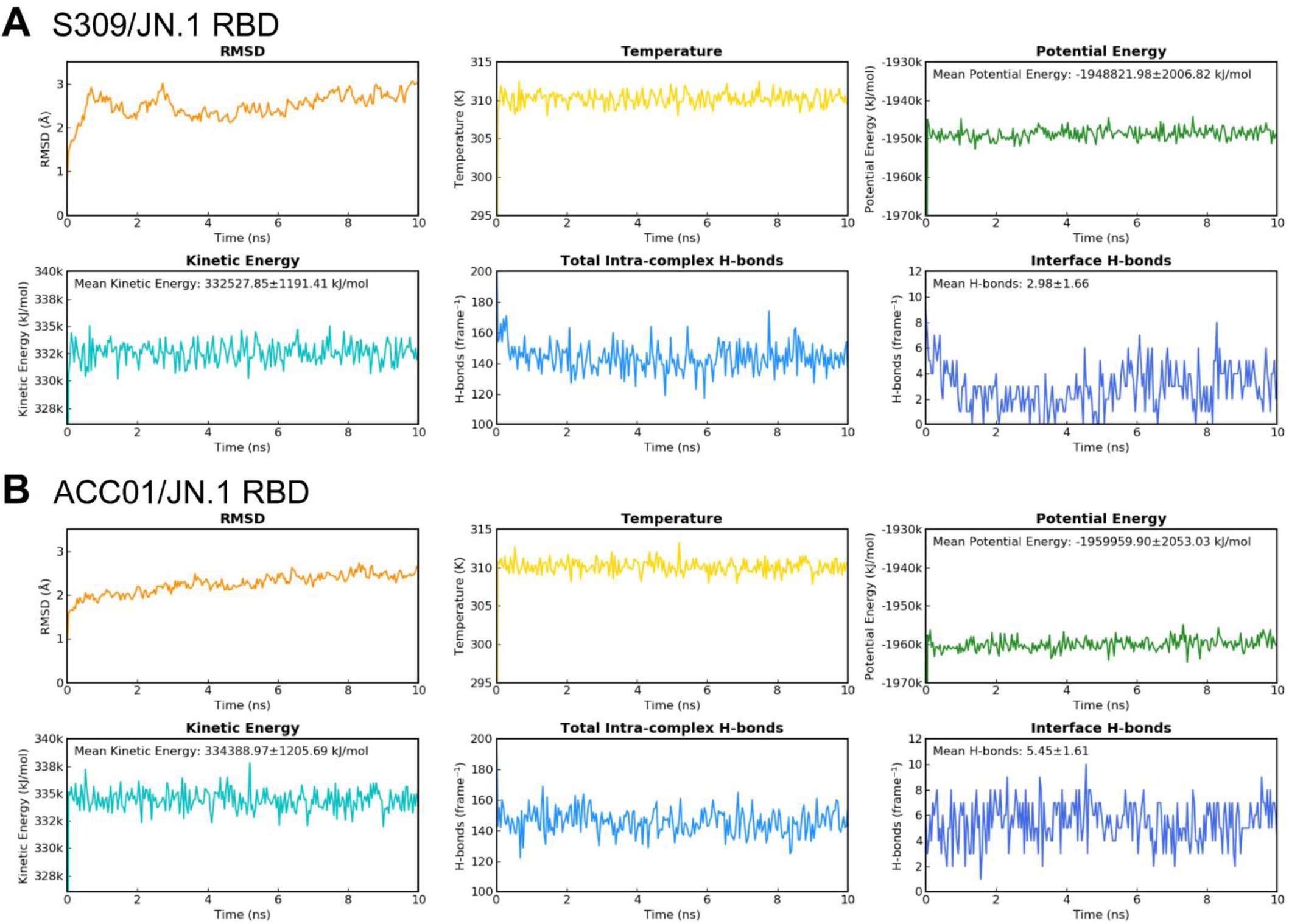
10-ns molecular dynamics (MD) simulation metrics for the S309/JN.1 RBD and ACC01/JN.1 RBD complexes (pure protein-protein interactions, Fig. 3). Time-evolution profiles are shown for: root-mean-square deviation (RMSD), temperature, potential energy, kinetic energy, total intra-complex hydrogen bonds, and specific antibody–antigen interface hydrogen bonds.

**Extended Data Figure 4.**
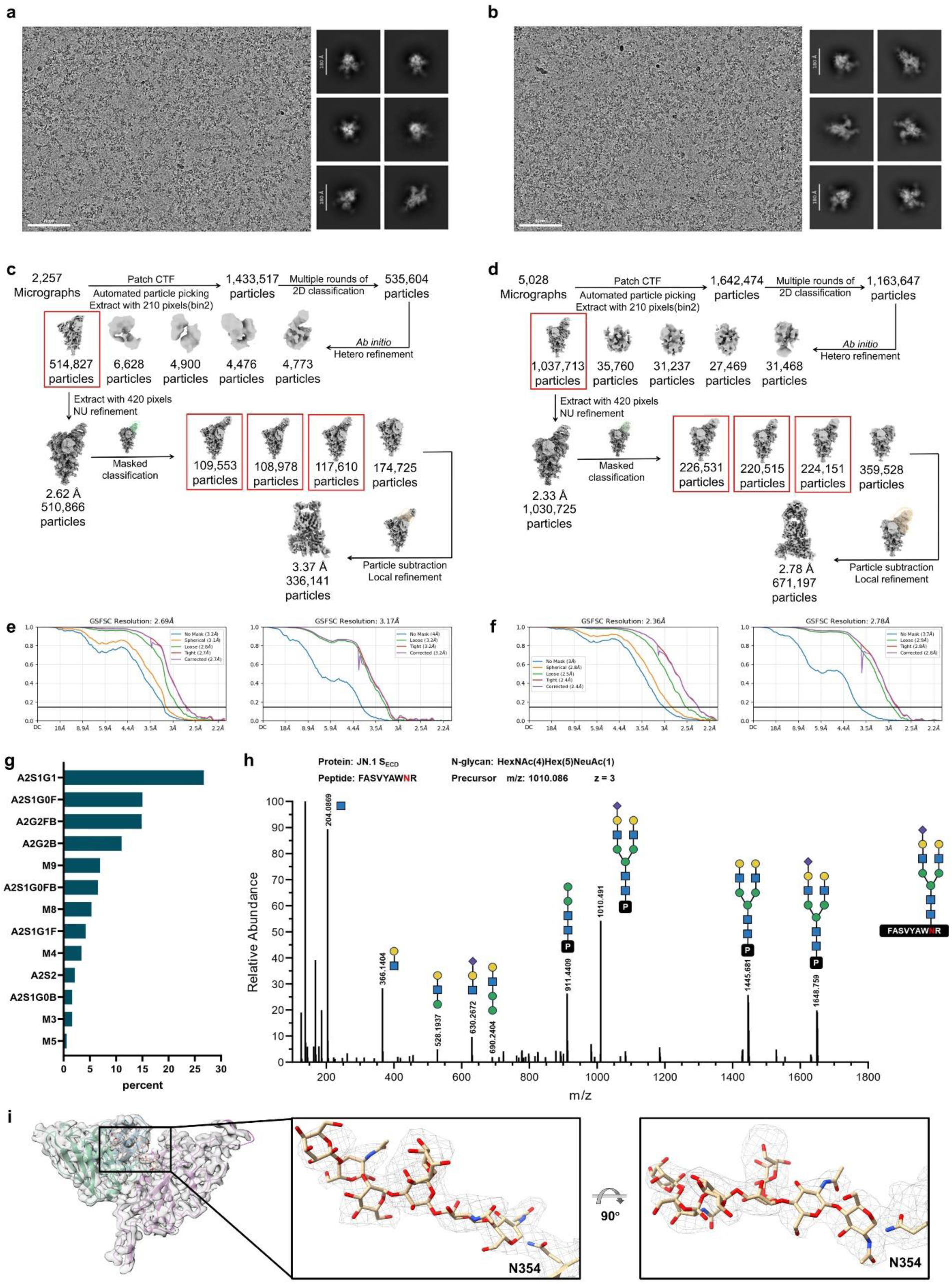
Cryo-EM single-particle analysis of antibody-RBD complexes and mass spectrometric characterization of the N354-linked glycan. **(a, b)** Representative electron micrograph and 2D class averages of **(a)** S309/JN.1 complex and **(b)** ACC01/JN.1 complex; **(c, d)** Data processing workflow, from purified protein samples to the final 3D reconstruction of **(c)** S309/JN.1 complex and **(d)** ACC01/JN.1 complex; **(e, f)** The Fourier Shell Correlation (FSC) curve of **(e)** S309/JN.1 complex and **(f)** ACC01/JN.1 complex. **(g)** Liquid chromatography-tandem mass spectrometry (LC-MS/MS) analysis revealing multiple N-glycan populations at Asn354. The predominant glycoform corresponds to HexNAc₄Hex₅NeuAc₁ (A2S1G1), with additional species annotated with their respective abundances; **(h)** Liquid chromatography-tandem mass spectrometry (LC-MS/MS) of the most abundant glycopeptides, with the predominant species corresponding to HexNAc₄Hex₅NeuAc₁ (theoretical *m/z* 1010.086, z=3; peptide FASVYAWNR), as shown in the main text (Fig. 4b); **(i)** Fitted model of ACC01/JN.1 complex presented as a ribbon diagram, with the cryo-EM map shown as transparent volume. Zoomed-in views of the glycan within the rectangle in the right panel are shown. The ordered core heptasaccharide accommodates the three most abundant glycoforms, collectively accounting for ∼60% of the N354 glycan population. The continuous density permits unambiguous tracing of the heptasaccharide core (Galβ1–4GlcNAcβ1–2Manα1–3(Manα1–6)Manβ1–4GlcNAcβ1–4GlcNAcβ1–N-Asn354) linked to Asn354. The well-resolved density supports the reliability of the atomic model presented in Fig. 4c.

**Extended Data Table 4.** Glycosidic torsion angles of the N354 glycan in the ACC01/JN.1 RBD complex. All measured φ, ψ, and ω torsion angles for the N354 glycan linkages fall within the expected low-energy conformational ranges derived from GlycoMapsDB and previous analyses of N-glycans. Linkages evaluated include Galβ1–4GlcNAc, GlcNAcβ1–2Man, Manα1–3Man, Manβ1–4GlcNAc, GlcNAcβ1–4GlcNAc, GlcNAcβ1–N-Asn, and the α1-6 branch Manα1–6Man. The glycan is presented from the non-reducing end (Gal) to the reducing end (Asn). Torsion angles were defined as described in the Methods. Briefly, φ and ψ follow the C+1 system (φ = O5–C1–O–Cx, ψ = C1–O–Cx–C(x+1)). For the Asn linkage, the φ angle is represented by the NAG1 acetamido torsion (C2–N2–C7–O7). For the α1-6 branch, ω = O6–C6–C5–C4. Status indicates the relative deviation from the expected range center, calculated as |*value* − *center*| / (*ℎalf* − *widt*ℎ) : Most Favored (deviation ≤ 0.33), Moderately Favored (0.33–0.67), Marginally Favored (0.67–1.0), and Disfavored (outside expected range). Marginally Favored and Disfavored categories were not observed. All measured values reside within favored regions, confirming that the glycan adopts a stable, unstrained conformation upon antibody binding. Abbreviations: GlcNAc, N-acetylglucosamine (NAG); Man, mannose (MAN/BMA); Gal, galactose (GAL); Asn, asparagine.

| Linkage<sup>1</sup> | Glycosidic bond | Torsion angle | Value (°) | Ideal center value (°) | Expected range (°)<sup>2</sup> | Status |
| --- | --- | --- | --- | --- | --- | --- |
| GAL6→NAG5 | Galβ1–4GlcNAc | $\phi$ | 107.0 | 110 | 80 to 140 | Most Favored |
| | | $\psi$ | –49.4 | –60 | –80 to –40 | Moderately Favored |
| NAG5→MAN4 | GlcNAcβ1–2Man | $\phi$ | 44.1 | 50 | 30 to 70 | Most Favored |
| | | $\psi$ | 92.4 | 95 | 70 to 120 | Most Favored |
| MAN4→BMA3 | Manα1–3Man | $\phi$ | –130.9 | –130 | –160 to –100 | Most Favored |
| | | $\psi$ | 81.8 | 80 | 60 to 100 | Most Favored |
| BMA3→NAG2 | Manβ1–4GlcNAc | $\phi$ | 116.7 | 110 | 80 to 140 | Most Favored |
| | | $\psi$ | –37.4 | –40 | –60 to –20 | Most Favored |
| NAG2→NAG1 | GlcNAcβ1–4GlcNAc | $\phi$ | 127.2 | 110 | 80 to 140 | Moderately Favored |
| | | $\psi$ | –69.4 | –60 | –80 to –40 | Moderately Favored |
| NAG1→Asn354 | GlcNAcβ1–N-Asn | $\phi$ | 179.8 | 180 | 120 to 240 | Most Favored |
| MAN8 → BMA3<br>(α1-6 branch) | Manα1–6Man | $\phi$ | 110.6 | 110 | 80 to 140 | Most Favored |
| | | $\psi$ | 123.1 | 130 | 100 to 160 | Most Favored |
| | | $\omega$ | –26.1 | 0 | –60 to 60 | Moderately Favored |

<sup>1</sup> All linkages are written in standard non-reducing to reducing end order.
<sup>2</sup> Expected ranges are derived from the GlycoMapsDB and literature values for common N-glycan linkages.

**Extended Data Figure 5.**
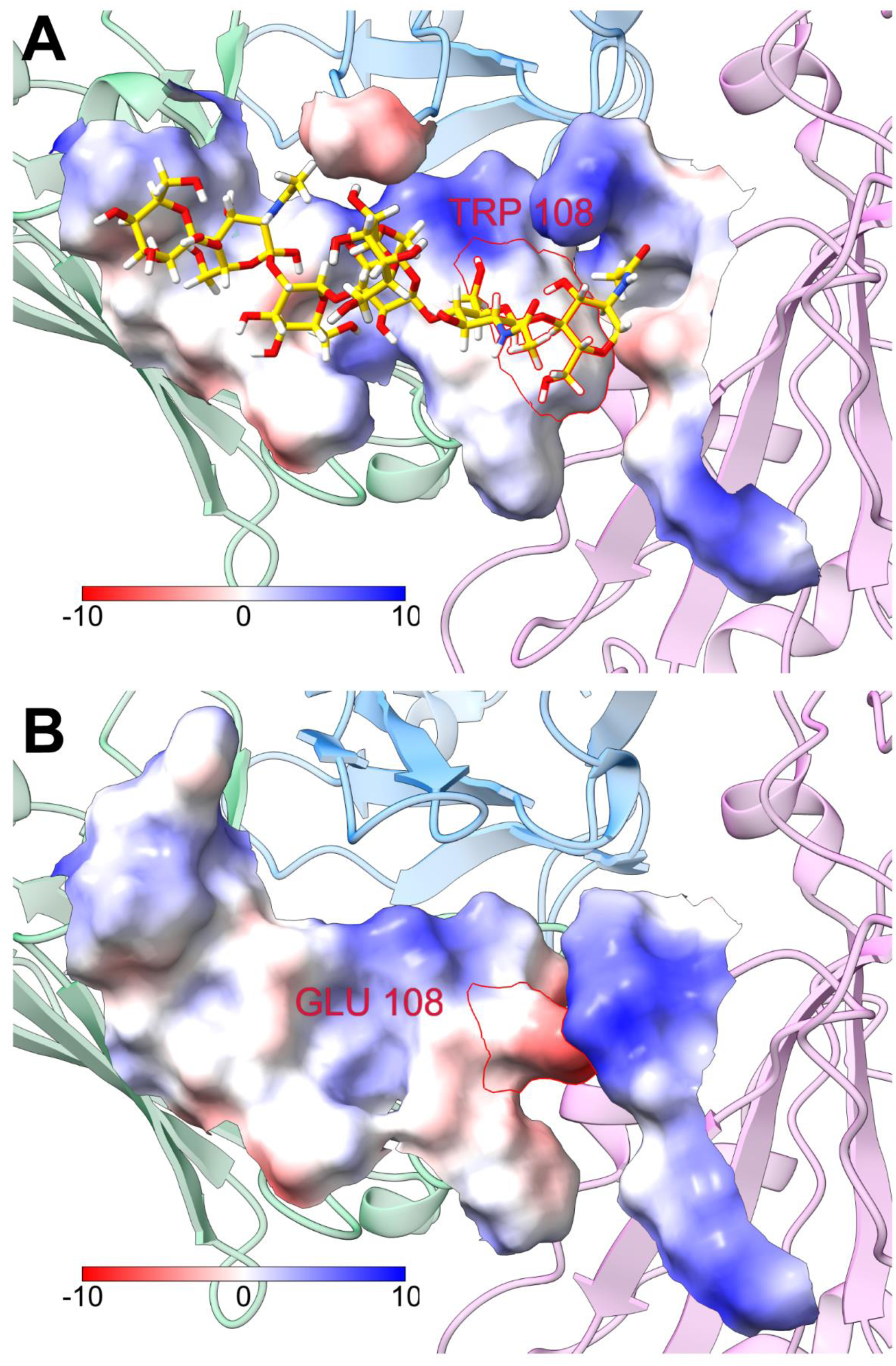
Electrostatic characteristics of the 5-Å glycan pockets in antibody-RBD complexes. **(a)** Visualization of the 5-Å pocket surrounding the N354-linked glycan in the ACC01/JN.1 RBD complex. The pocket is rendered as electrostatic potential surface, while the rest of the protein is shown in cartoon mode and colored by individual chains (heavy chain: green; light chain: blue; RBD: orchid). The residue E108W (red) abuts the glycan core, and this local region exhibits electrostatic neutrality. **(b)** Visualization of the pseudo-glycan 5-Å pocket in the S309/JN.1 RBD complex. Since S309 cannot stabilize the N354-linked glycan, the S309/JN.1 complex was structurally aligned to the ACC01/JN.1 complex, and the pseudo-glycan pocket was constructed based on the glycan position in ACC01/JN.1. The surface and cartoon representation as well as chain coloring are consistent with those in panel **(a)**. The corresponding glutamate residue (red) abuts the putative glycan core and confers local electronegativity. Given that the root region (NAG1, NAG2, BMA3) of the complex-type N-glycan at N354 is intrinsically electrostatically neutral, the electrostatic potential alteration at residue 108 is unlikely to exert a substantial impact on glycan stabilization.

**Extended Data Table 5.** Atomic contacts between the N354 glycan (chain F) and the ACC01 antibody (chains A and B) or JN.1 RBD antigen (chain M). Contacts were calculated using UCSF ChimeraX contacts command with default parameters (overlap cutoff –0.4). The table lists all detected atom pairs. Summary: chain A (antibody heavy chain) participates in 47 contacts (92.2%), chain B (antibody light chain) in 1 contact (2.0%), chain M (JN.1 RBD antigen) in 3 contacts (5.9%).

| # | Glycan atom | Protein atom | Distance (Å) | Overlap |
| --- | --- | --- | --- | --- |
| 1 | /F NAG 5 H1 | /A ASN 59 OD1 | 1.708 | 0.372 |
| 2 | /F NAG 5 O1 | /A ASN 59 OD1 | 2.23 | 0.35 |
| 3 | /F NAG 2 H61 | /A TRP 108 HB2 | 1.663 | 0.337 |
| 4 | /F NAG 1 H81 | /M ARG 346 O | 2.198 | 0.282 |
| 5 | /F NAG 5 O1 | /A ASN 59 CG | 2.933 | 0.267 |
| 6 | /F NAG 2 C6 | /A TRP 108 HB2 | 2.571 | 0.129 |
| 7 | /F MAN 4 C6 | /A ASN 57 HB3 | 2.605 | 0.095 |
| 8 | /F MAN 4 H61 | /A ASN 57 ND2 | 2.554 | 0.071 |
| 9 | /F NAG 5 O6 | /A TYR 60 H | 2.034 | 0.066 |
| 10 | /F NAG 2 H61 | /A TRP 108 CB | 2.642 | 0.058 |
| 11 | /F MAN 4 H4 | /A THR 58 O | 2.482 | -0.002 |
| 12 | /F NAG 5 H1 | /A ASN 59 CG | 2.73 | -0.03 |
| 13 | /F MAN 4 H61 | /A ASN 57 HB3 | 2.035 | -0.035 |
| 14 | /F NAG 1 C8 | /M ARG 346 O | 3.227 | -0.047 |
| 15 | /F NAG 5 C1 | /A ASN 59 OD1 | 3.242 | -0.062 |
| 16 | /F NAG 5 HO6 | /A GLN 65 OE1 | 2.161 | -0.081 |
| 17 | /F NAG 2 C6 | /A GLY 107 O | 3.282 | -0.102 |
| 18 | /F NAG 2 C6 | /A TRP 108 CB | 3.503 | -0.103 |
| 19 | /F NAG 5 H61 | /A THR 58 HG1 | 2.108 | -0.108 |
| 20 | /F NAG 5 HO3 | /A GLN 65 HE22 | 2.132 | -0.132 |
| 21 | /F NAG 2 H62 | /A GLY 107 O | 2.613 | -0.133 |
| 22 | /F MAN 4 H61 | /A ASN 57 CB | 2.839 | -0.139 |
| 23 | /F NAG 5 O5 | /A ASN 59 OD1 | 3.15 | -0.17 |
| 24 | /F MAN 4 H61 | /A ASN 57 HD21 | 2.173 | -0.173 |
| 25 | /F MAN 4 H61 | /A ASN 57 CG | 2.881 | -0.181 |
| 26 | /F NAG 5 O5 | /A ASN 59 HA | 2.688 | -0.188 |
| 27 | /F MAN 4 C6 | /A ASN 57 CB | 3.599 | -0.199 |
| 28 | /F NAG 5 O6 | /A TYR 60 CD2 | 3.403 | -0.203 |
| 29 | /F NAG 5 H61 | /A THR 58 O | 2.689 | -0.209 |
| 30 | /F NAG 5 O5 | /A THR 58 O | 3.209 | -0.229 |
| 31 | /F NAG 5 O6 | /A TYR 60 CB | 3.439 | -0.239 |
| 32 | /F NAG 5 O6 | /A TYR 60 N | 2.97 | -0.245 |
| 33 | /F MAN 4 C4 | /A THR 58 O | 3.442 | -0.262 |
| 34 | /F NAG 5 O6 | /A TYR 60 HB3 | 2.768 | -0.268 |
| 35 | /F NAG 2 O6 | /A TRP 108 HB2 | 2.779 | -0.279 |
| 36 | /F MAN 4 O5 | /A ASN 59 OD1 | 3.263 | -0.283 |
| 37 | /F NAG 2 O6 | /A TRP 108 CB | 3.5 | -0.3 |
| 38 | /F NAG 2 H61 | /A TRP 108 CA | 3.002 | -0.302 |
| 39 | /F MAN 4 C6 | /A ASN 57 ND2 | 3.637 | -0.312 |
| 40 | /F NAG 5 O3 | /A GLN 65 HE22 | 2.427 | -0.327 |
| 41 | /F MAN 4 H62 | /A ASN 57 HB3 | 2.329 | -0.329 |
| 42 | /F NAG 2 C6 | /A TRP 108 CA | 3.74 | -0.34 |
| 43 | /F MAN 4 H62 | /A THR 58 O | 2.824 | -0.344 |
| 44 | /F NAG 2 C6 | /A GLY 107 C | 3.748 | -0.348 |
| 45 | /F NAG 5 C6 | /A THR 58 HG1 | 3.057 | -0.357 |
| 46 | /F NAG 1 O7 | /M ARG 346 O | 3.317 | -0.357 |
| 47 | /F NAG 5 O7 | /B THR 94 CG2 | 3.555 | -0.375 |
| 48 | /F NAG 5 O6 | /A TYR 60 CG | 3.576 | -0.376 |
| 49 | /F NAG 5 HO3 | /A TYR 60 O | 2.458 | -0.378 |
| 50 | /F GAL 6 H5 | /A GLN 65 CG | 3.093 | -0.393 |
| 51 | /F NAG 5 HO6 | /A GLN 65 CD | 3.1 | -0.4 |

\*Chain identification: A = antibody heavy chain, B = antibody light chain, M = JN.1 RBD antigen, F = N354 glycan. NAG = GlcNAc, MAN = mannose (including BMA), GAL = galactose.\*

## Reference

1. Li D, Wang L, Maziuk BF, Yao X, Wolozin B, Cho YK. Directed evolution of a picomolar-affinity, high-specificity antibody targeting phosphorylated tau. The Journal of biological chemistry. 2018;293(31):12081–94.

2. Liu P, Yue C, Meng B, Xiao T, Yang S, Liu S, et al. Spike N354 glycosylation augments SARS-CoV-2 fitness for human adaptation through structural plasticity. National science review. 2024;11(7):nwae206.

3. Li P, Faraone JN, Hsu CC, Chamblee M, Zheng YM, Carlin C, et al. Neutralization escape, infectivity, and membrane fusion of JN.1-derived SARS-CoV-2 SLip, FLiRT, and KP.2 variants. Cell reports. 2024;43(8):114520.

4. Contractor D, Globisch C, Swaroop S, Jain A. Structural basis of Omicron immune evasion: A comparative computational study. Computers in biology and medicine. 2022;147:105758.

5. Steiner S, Kratzel A, Barut GT, Lang RM, Aguiar Moreira E, Thomann L, et al. SARS-CoV-2 biology and host interactions. Nature reviews Microbiology. 2024;22(4):206–25.

6. Wang Q, Iketani S, Li Z, Liu L, Guo Y, Huang Y, et al. Alarming antibody evasion properties of rising SARS-CoV-2 BQ and XBB subvariants. Cell. 2023;186(2):279–86.e8.

7. Yajima H, Anraku Y, Kaku Y, Kimura KT, Plianchaisuk A, Okumura K, et al. Structural basis for receptor-binding domain mobility of the spike in SARS-CoV-2 BA.2.86 and JN.1. Nature communications. 2024;15(1):8574.

8. Chi X, Xia L, Zhang G, Chi X, Huang B, Zhang Y, et al. Comprehensive structural analysis reveals broad-spectrum neutralizing antibodies against SARS-CoV-2 Omicron variants. Cell discovery. 2023;9(1):37.

9. Pinto D, Park YJ, Beltramello M, Walls AC, Tortorici MA, Bianchi S, et al. Cross-neutralization of SARS-CoV-2 by a human monoclonal SARS-CoV antibody. Nature. 2020;583(7815):290–5.

10. Afzal M, Melnyk D, Courty T, Schimanski L, Hill M, Neil S, et al. Determination of resilience of a panel of broadly neutralizing mAbs to emerging variants of SARS-CoV-2 generated using reverse genetics. iScience. 2025;28(6):112451.

11. Simons JF, Lim YW, Carter KP, Wagner EK, Wayham N, Adler AS, et al. Affinity maturation of antibodies by combinatorial codon mutagenesis versus error-prone PCR. mAbs. 2020;12(1):1803646.

12. Arnold FH. Directed evolution: bringing new chemistry to life. Angewandte Chemie (International ed in English). 2017;57(16):4143.

13. Wellner A, McMahon C, Gilman MSA, Clements JR, Clark S, Nguyen KM, et al. Rapid generation of potent antibodies by autonomous hypermutation in yeast. Nature chemical biology. 2021;17(10):1057–64.

14. Bai G, Sun C, Guo Z, Wang Y, Zeng X, Su Y, et al. Accelerating antibody discovery and design with artificial intelligence: Recent advances and prospects. Seminars in cancer biology. 2023;95:13–24.

15. Hie B, Zhong ED, Berger B, Bryson B. Learning the language of viral evolution and escape. Science (New York, NY). 2021;371(6526):284–8.

16. Hie BL, Shanker VR, Xu D, Bruun TUJ, Weidenbacher PA, Tang S, et al. Efficient evolution of human antibodies from general protein language models. Nature biotechnology. 2024;42(2):275–83.

17. Zhang Q, Chen W, Qin M, Wang Y, Pu Z, Ding K, et al. Integrating protein language models and automatic biofoundry for enhanced protein evolution. Nature communications. 2025;16(1):1553.

18. Nijkamp E, Ruffolo JA, Weinstein EN, Naik N, Madani A. ProGen2: Exploring the boundaries of protein language models. Cell systems. 2023;14(11):968–78.e3.

19. Desautels TA, Arrildt KT, Zemla AT, Lau EY, Zhu F, Ricci D, et al. Computationally restoring the potency of a clinical antibody against Omicron. Nature. 2024;629(8013):878–85.

20. Fernandez-de-Cossio-Diaz J, Uguzzoni G, Ricard K, Anselmi F, Nizak C, Pagnani A, et al. Inference and design of antibody specificity: From experiments to models and back. PLoS computational biology. 2024;20(10):e1012522.

21. Zhou L, Pacchiardi L, Martínez-Plumed F, Collins KM, Moros-Daval Y, Zhang S, et al. General scales unlock AI evaluation with explanatory and predictive power. Nature. 2026;652(8108):58–67.

22. Wang B, Gallolu Kankanamalage S, Dong J, Liu Y. Optimization of therapeutic antibodies. Antibody therapeutics. 2021;4(1):45–54.

23. Sela-Culang I, Kunik V, Ofran Y. The structural basis of antibody-antigen recognition. Frontiers in immunology. 2013;4:302.

24. Syrgkanis V, Lewis G, Oprescu M, Hei M, Battocchi K, Dillon E, et al. Causal Inference and Machine Learning in Practice with EconML and CausalML: Industrial Use Cases at Microsoft, TripAdvisor, Uber. Proceedings of the 27th ACM SIGKDD Conference on Knowledge Discovery & Data Mining; Virtual Event, Singapore: Association for Computing Machinery; 2021. p. 4072–3.

25. Martí D, Alsina M, Alemán C, Bertran O, Turon P, Torras J. Unravelling the molecular interactions between the SARS-CoV-2 RBD spike protein and various specific monoclonal antibodies. Biochimie. 2022;193:90–102.

26. Cummings RD, Schnaar RL, Esko JD, Woods RJ, Drickamer K, Taylor ME. Principles of Glycan Recognition. In: Varki A, Cummings RD, Esko JD, Stanley P, Hart GW, Aebi M, et al., editors. Essentials of Glycobiology. Cold Spring Harbor (NY): Cold Spring Harbor Laboratory Press Copyright © 2022 The Consortium of Glycobiology Editors, La Jolla, California; published by Cold Spring Harbor Laboratory Press; doi:10.1101/glycobiology.4e.29. All rights reserved.; 2022. p. 387–402.

27. Böhm M, Bohne-Lang A, Frank M, Loss A, Rojas-Macias MA, Lütteke T. Glycosciences. DB: an annotated data collection linking glycomics and proteomics data (2018 update). Nucleic acids research. 2019;47(D1):D1195–d201.

28. Allen JD, Ivory DP, Song SG, He WT, Capozzola T, Yong P, et al. The diversity of the glycan shield of sarbecoviruses related to SARS-CoV-2. Cell reports. 2023;42(4):112307.

29. Mellis IA, Wu M, Hong H, Tzang CC, Bowen A, Wang Q, et al. Antibody evasion and receptor binding of SARS-CoV-2 LP.8.1.1, NB.1.8.1, XFG, and related subvariants. Cell reports. 2025;44(10):116440.

30. Cao G, Xu C, Wang L, Chai K, Wu B. Global Surveillance and Biological Characterization of the SARS-CoV-2 NB.1.8.1 Variant: An Emerging VUM Lineage Under Scrutiny. Viruses. 2025;17(11).

31. Cao Y, Song W, Wang L, Liu P, Yue C, Jian F, et al. Characterization of the enhanced infectivity and antibody evasion of Omicron BA.2.75. Cell host & microbe. 2022;30(11):1527–39.e5.

32. Meng EC, Goddard TD, Pettersen EF, Couch GS, Pearson ZJ, Morris JH, et al. UCSF ChimeraX: Tools for structure building and analysis. Protein science : a publication of the Protein Society. 2023;32(11):e4792.

33. Schymkowitz J, Borg J, Stricher F, Nys R, Rousseau F, Serrano L. The FoldX web server: an online force field. Nucleic acids research. 2005;33(Web Server issue):W382–8.

34. Chernozhukov V, Chetverikov D, Demirer M, Duflo E, Hansen C, Newey W, et al. Double/debiased machine learning for treatment and structural parameters. The Econometrics Journal. 2018;21(1):C1–C68.

35. Schrodinger, LLC. The PyMOL Molecular Graphics System, Version 1.8. 2015.

36. Eastman P, Galvelis R, Peláez RP, Abreu CRA, Farr SE, Gallicchio E, et al. OpenMM 8: Molecular Dynamics Simulation with Machine Learning Potentials. The journal of physical chemistry B. 2024;128(1):109–16.

37. Case DA, Aktulga HM, Belfon K, Cerutti DS, Cisneros GA, Cruzeiro VWD, et al. AmberTools. Journal of chemical information and modeling. 2023;63(20):6183–91.

38. Salomon-Ferrer R, Götz AW, Poole D, Le Grand S, Walker RC. Routine Microsecond Molecular Dynamics Simulations with AMBER on GPUs. 2. Explicit Solvent Particle Mesh Ewald. Journal of chemical theory and computation. 2013;9(9):3878–88.

39. Nayar D, Agarwal M, Chakravarty C. Comparison of Tetrahedral Order, Liquid State Anomalies, and Hydration Behavior of mTIP3P and TIP4P Water Models. Journal of chemical theory and computation. 2011;7(10):3354–67.

40. Maaten Lvd, Hinton GE. Visualizing Data using t-SNE. Journal of Machine Learning Research. 2008;9:2579–605.

41. McInnes L, Healy J. UMAP: Uniform Manifold Approximation and Projection for Dimension Reduction. 2018.

42. Tenenbaum JB, de Silva V, Langford JC. A global geometric framework for nonlinear dimensionality reduction. Science (New York, NY). 2000;290(5500):2319–23.

43. Cope D, Blakeslee B, McCourt ME. Analysis of multidimensional difference-of-Gaussians filters in terms of directly observable parameters. Journal of the Optical Society of America A, Optics, image science, and vision. 2013;30(5):1002–12.

44. Stone JE, Vandivort KL, Schulten K, editors. GPU-accelerated molecular visualization on petascale supercomputing platforms. Proceedings of the 8th International Workshop on Ultrascale Visualization; 2013.

45. Humphrey W, Dalke A, Schulten K. VMD: visual molecular dynamics. Journal of molecular graphics. 1996;14(1):33–8, 27-8.

46. Qu L, Qiao X, Qi F, Nishida N, Hoshino T. Analysis of Binding Modes of Antigen-Antibody Complexes by Molecular Mechanics Calculation. Journal of chemical information and modeling. 2021;61(5):2396–406.

47. Donald JE, Kulp DW, DeGrado WF. Salt bridges: geometrically specific, designable interactions. Proteins. 2011;79(3):898–915.

48. Land H, Humble MS. YASARA: A Tool to Obtain Structural Guidance in Biocatalytic Investigations. Methods in molecular biology (Clifton, NJ). 2018;1685:43–67.

49. Katoh K, Standley DM. MAFFT multiple sequence alignment software version 7: improvements in performance and usability. Molecular biology and evolution. 2013;30(4):772–80.

50. Minh BQ, Schmidt HA, Chernomor O, Schrempf D, Woodhams MD, von Haeseler A, et al. IQ-TREE 2: New Models and Efficient Methods for Phylogenetic Inference in the Genomic Era. Molecular biology and evolution. 2020;37(5):1530–4

